# Evaluating the dependence of ADC-fMRI on haemodynamics in breath-hold and resting-state conditions

**DOI:** 10.1101/2025.05.15.654237

**Authors:** Inès de Riedmatten, Arthur P C Spencer, Jasmine Nguyen-Duc, Jean-Baptiste Pérot, Filip Szczepankiewicz, Oscar Esteban, Ileana O Jelescu

## Abstract

Apparent diffusion coefficient (ADC)-fMRI offers a promising functional contrast, capable of mapping neuronal activity directly in both grey and white matter. However, previous studies have shown that diffusion-weighted fMRI (dfMRI), from which ADC-fMRI derives, is influenced by BOLD effects, leading to a concern that the dfMRI contrast is still rooted in neurovascular rather than neuromorphological coupling. Mitigation strategies have been proposed to remove vascular contributions while retaining neuromorphological coupling, by: i) analysing ADC timecourses calculated from two interleaved diffusion-weightings, known as ADC-fMRI; ii) using b-values of at least 200 s mm^−2^; and iii) using a sequence compensated for cross-terms with fluctuating background field gradients associated with blood oxygenation. Respiration-induced haemodynamic fluctuations, which are dissociated from neural activity, are an excellent test-bed for the robustness of ADC-fMRI to vascular contributions. In this study, we investigate the association between end-tidal CO_2_ and ADC-fMRI, in comparison with dfMRI and BOLD, in both breath-hold and resting-state paradigms in the human brain. We confirm a strong dependence of the BOLD signal on respiration, and a pattern of delayed haemodynamic response in white matter. While dfMRI mitigates much of the vascular contribution, it retains some association with respiration, as expected. Conversely, ADC-fMRI is mostly unaffected by vascular contribution, exhibiting minimal correlation between expired CO_2_ and ADC timeseries, as well as low inter- and intra-subject reproducibility in correlation maps. These findings validate ADC-fMRI as a predominantly non-vascular contrast sensitive to microstructural dynamics, enabling whole-brain functional imaging unconstrained by vascular confounds.

## 1 Introduction

Functional MRI (fMRI) methods for probing neuronal activity non-invasively continue to push the boundaries of spatial and temporal resolution, yet blood-oxygen level dependent (BOLD)-fMRI remains inherently constrained by the indirect nature of neurovascular coupling (Logothetis et al., 2001; O’Herron et al., 2016). This contrast links neuronal activity to fluctuations in the ratio of oxy-to deoxyhaemoglobin within blood vessels via the haemodynamic response. Conversely, apparent diffusion coefficient (ADC)-fMRI offers a fundamentally different approach, capturing the microstructural dynamics associated with neuronal activity, independent of haemodynamic confounds (Abe et al., 2017, 2020; Darquié et al., 2001; Komaki et al., 2020; Le Bihan et al., 2006; Olszowy et al., 2021; Tsurugizawa et al., 2013). By leveraging neuromorphological rather than neurovascular coupling, ADC-fMRI holds the promise of a more direct window into brain function. More specifically, the ADC-fMRI contrast is sensitive to cellular deformations triggered by neuronal activity. These morphological changes occur in neurites (Ling et al., 2020), synaptic boutons (Chéreau et al., 2017), myelinated axons (Kwon et al., 2023), neuronal somas (Ling et al., 2020), and astrocytic processes (Sherpa et al., 2016), altering the diffusion of water molecules and producing detectable variations in the ADC timeseries (Abe et al., 2017; Darquié et al., 2001; Nunes et al., 2021). Previous studies have demonstrated that ADC is a valuable contrast to detect excitatory neuronal activity in task-based fMRI (Darquié et al., 2001; Nguyen-Duc et al., 2025; Nicolas et al., 2017; Nunes et al., 2021; Olszowy et al., 2021; Spees et al., 2013; Spencer et al., 2025), and to identify resting-state networks (de Riedmatten et al., 2025; Olszowy et al., 2021). The raw diffusion-weighted timeseries was initially proposed as a functional contrast in its own right, referred to as diffusion fMRI (dfMRI). However, the dfMRI signal is intrinsically influenced by the BOLD contrast via T_2_-weighting which likely supersedes its sensitivity to microstructure fluctuations (K. L. Miller et al., 2007). While most T_2_-weighting is removed from ADC timecourses by calculating a ratio of consecutive signals, an ongoing debate remains regarding whether ADC-fMRI truly provides a neuromorphological contrast independent of vascular contributions. In this study, we address this question by eliciting a non-neuronal BOLD response through breath-hold tasks and examining whether this response appears in the ADC-fMRI timeseries. Additionally, we assess the extent to which ADC-fMRI and BOLD are each influenced by spontaneous fluctuations in resting breathing patterns during resting-state acquisitions.

### 1.1 BOLD Signal and Respiration

During neuronal activity, the local blood flow and volume increase to meet and eventually exceed the increased oxygen metabolic demand, increasing the oxy-vs deoxyhaemoglobin ratio. As deoxyhaemoglobin is paramagnetic, neuronal activity is captured as an increase in the T_2_ (or 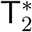) BOLD signal. Notably, a similar BOLD signal increase can be elicited in the absence of neuronal activity by voluntarily altering the arterial CO_2_ level (Birn, Smith, et al., 2008; Chang et al., 2008). As CO_2_ is a potent vasodilator, an increase in arterial blood CO_2_ triggers cerebral vasodilation, leading to increased cerebral blood flow, increased venous oxy-to deoxyhaemoglobin ratio, and consequently increased BOLD signal (Addeh et al., 2025; Birn et al., 2006; Chang & Glover, 2009).

A simple way to induce hypercapnia is with a breath-hold task (Pinto et al., 2021). During the task, fluctuations in respiration depth and rate can be monitored using a gas analyser (Addeh et al., 2025; Birn et al., 2006; Wise et al., 2004). Notably, the time-lagged end-tidal CO_2_ (p_ET_CO_2_) has been shown to strongly correlate with the BOLD signal (Murphy et al., 2011; Zvolanek et al., 2023). This lag reflects both the transit time of blood from the lungs to the brain, and the local vasoreactivity delays, the latter of which has been shown to contribute the most (Chang et al., 2008; Pinto et al., 2021; Wise et al., 2004). The lags vary across the brain by several seconds, likely due to regional heterogeneity in the vascular response (Birn, Murphy, & Bandettini, 2008; Gong et al., 2023). Breath-hold tasks have been widely used to demonstrate the confounding effects of respiration on BOLD-fMRI analysis (Chang et al., 2008), in addition to deriving cerebrovascular reactivity (CVR) maps, which represent the blood flow response to a vasoactive stimulus (Zvolanek et al., 2023). Given their well-established impact on vascular signals, breath-hold tasks therefore serve as a valuable tool for assessing the extent to which dfMRI and ADC-fMRI contrasts are influenced by vascular contributions.

The breath-hold task elicits a robust BOLD response due to the substantial accumulation of CO_2_ in the blood. While this response provides valuable insight into the vascular contributions to the MRI signal, it does not necessarily reflect typical physiological conditions outside the breath-hold paradigm, since resting breathing results in smaller fluctuations in blood CO_2_ levels. Nevertheless, even spontaneous p_ET_CO_2_ variations have been shown to correlate significantly with the BOLD signal during resting-state experiments (Chang & Glover, 2009; Wise et al., 2004). Specifically, increased respiration depth or rate have been associated with a decrease in BOLD-fMRI signal (Birn, Smith, et al., 2008). Regions exhibiting a strong association between p_ET_CO_2_ and BOLD-fMRI largely coincide with the default mode network regions (Birn et al., 2006; Chang et al., 2009). Given that the frequency range of these signals is similar (*<* 0.1 Hz for BOLD and 0-0.05 Hz for p_ET_CO_2_, in resting-state), they cannot be easily separated from each other, and fluctuations in the breathing pattern may introduce spurious functional connectivity (Birn et al., 2006; Chang et al., 2009; Wise et al., 2004).

### 1.2 ADC-fMRI and Vascular Contamination

In an ADC-fMRI acquisition, the ADC at a given time point is calculated from two diffusion-weighted signals 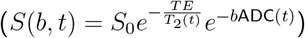 each acquired with different b-values:

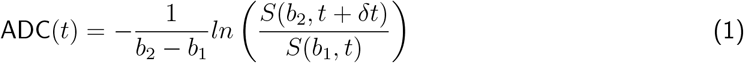

The diffusion-weighted MRI signal is influenced by haemodynamic fluctuations via three key mechanisms, each of which can be mitigated by careful design of the ADC-fMRI acquisition (Olszowy et al., 2021). Firstly, at low b-values, the diffusion signal is sensitive to fast-moving spins in capillaries, and to voxel-level blood volume changes (Le Bihan et al., 2006). This phenomenon is referred to as perfusion, or pseudo-diffusion in intra-voxel incoherent motion (IVIM) contrast. To limit the perfusion contribution, we selected b-values of at least 200 s mm^−2^, where perfusion accounts for around 1% of the measured diffusion signal and is considered negligible (Le Bihan, 2019). This approach of avoiding the perfusion regime in the ADC calculation has previously been described as ‘synthetic’ or ‘shifted’ ADC (Debaker et al., 2020; Iima & Le Bihan, 2016). Second, the diffusion-weighted signal *S*(*b, t*) is a function of T_2_, and therefore sensitive to the BOLD contrast. Taking the ratio between two diffusion-weighted signals as described in Equation 1 largely reduces this contribution. Finally, magnetic susceptibility-induced background field gradients (G_*s*_) vary around blood vessels during the haemodynamic response, and affect the effective b-value 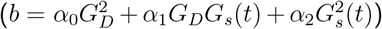, the raw diffusion-weighted signals, and the ADC (Pampel et al., 2010). We minimised these contributions with a cross-term compensated acquisition.

Previous studies mitigating all three mechanisms of vascular contamination have acquired ADC-fMRI with linear diffusion encoding (imparting diffusion weighting along a single arbitrary direction in space) with bipolar gradients for cross-term compensation (de Riedmatten et al., 2025; Nguyen-Duc et al., 2025; Olszowy et al., 2021). A more recent study used isotropic diffusion encoding, which imparts diffusion-weighting in all directions, and thus further provides sensitivity to white matter microstructure fluctuations independently of underlying fibre organisation (Spencer et al., 2025). Building on this work, we used a similar isotropic ADC-fMRI acquisition protocol, but minimised cross-terms with a compensated gradient waveform for isotropic diffusion encoding (Szczepankiewicz & Sjolund, 2021).

While previous studies have implemented measures to minimise some combination of these three contributions (De Luca et al., 2019; de Riedmatten et al., 2025; Nguyen-Duc et al., 2025; Nicolas et al., 2017; Olszowy et al., 2021; Spencer et al., 2025), they have not directly assessed the extent to which the resulting ADC-fMRI contrast is independent of haemodynamic fluctuations. In this study, we compared the BOLD-fMRI and the ADC-fMRI response to hypercapnia (with a breath-hold task) and normocapnia (with resting-state), while monitoring CO_2_ variations. This aims to reveal the extent to which the ADC-fMRI contrast is contaminated by BOLD effects.

## 2 Methods

### 2.1 Experiments

This study was approved by the ethics committee of the canton of Vaud, Switzerland (CER-VD). All participants provided written informed consent. Healthy participants (n = 15) underwent four breath-hold runs (two with ADC-fMRI and two with BOLD-fMRI) and two resting-state runs (one with ADC-fMRI and one with BOLD-fMRI), in one session. The order of breath-hold runs and resting-state runs was alternated from one subject to the next to avoid any systematic impact of accumulating hypercapnia in a specific contrast.

#### 2.1.1 Breath-hold

Visual cues for the breath-hold tasks were generated using PsychoPy (Peirce et al., 2019). Each run consisted of four repetitions of the task epoch, where one epoch comprised 18 s resting breathing, a 14 s breath-hold period, followed by 16 s resting breathing, for a total of 48 s per epoch and 192 s per complete breath-hold run. Resting breathing periods were self-paced, apart from a cued 2 s inhale and 2 s exhale immediately before the breath-hold period, and a 2 s exhale immediately after the breath-hold period, to ensure accurate p_ET_CO_2_ measurements (Pinto et al., 2021). The breath-hold paradigm is summarised in Figure 1A. Breathing was monitored via a nasal cannula, as described below.

**Figure 1.**
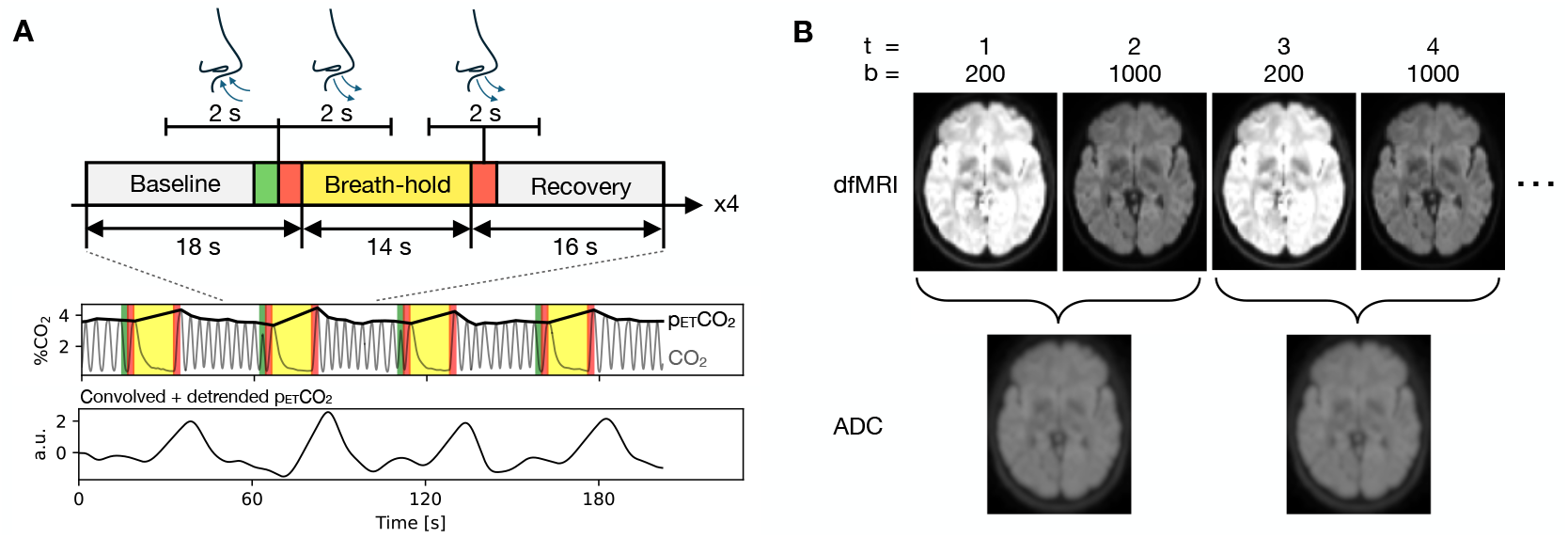
Breath-hold task paradigm, p_ET_CO_2_ acquisition and ADC-fMRI acquisition. A) The task consisted of four epochs of a 14 s breath-hold in between periods of self-paced resting breathing (baseline and recovery periods). An inhale and exhale were cued before holding, and an exhale after. CO_2_ was recorded with a gas analyser via a nasal cannula, from which p_ET_CO_2_ was measured by interpolating between peak CO_2_ at each exhale. This was convolved with the haemodynamic response function and detrended prior to correlation with fMRI data. B) The ADC-fMRI timeseries was calculated from interleaved b-value pairs with b = 200 and 1000 s mm^−2^. Each individual b-value dfMRI timeseries was also analysed.

#### 2.1.2 Resting-state

Resting-state runs lasted 14 minutes 40 s. Participants were instructed to stare at a fixation cross, not think about anything in particular, try not to fall asleep, and to breathe through their nose (to allow CO_2_ measurements via the nasal cannula).

### 2.2 MRI Acquisition

MRI data were acquired using a 3T Siemens Magnetom Prisma with 80 mT/m gradients and 200 T/m/s slew rate, and a 64-channel head coil. Whole-brain T_1_-weighted images were acquired, for anatomical reference and tissue segmentation, using 3D Magnetization Prepared 2 Rapid Acquisition Gradient Echoes (MP2RAGE) (Marques et al., 2010) with the following parameters: 1 mm^3^ isotropic voxels; 256 x 256 mm^2^ field of view; 176 slices; repetition time (TR) 5000 ms; echo time (TE) 2.98 ms; inversion times (TI) 700, 2500 ms; flip angles 4°, 5°; in-plane acceleration factor 3 using Generalized Autocalibrating Partially Parallel Acquisitions (GRAPPA) (Griswold et al., 2002).

DfMRI data were acquired with a diffusion-weighted spin-echo EPI sequence that enables spherical diffusion encoding, using a waveform compensated for cross-terms with background field gradients (Szczepankiewicz & Sjölund, 2021). The waveform was generated using the NOW toolbox (https://github.com/jsjol/NOW). The following parameters were used for the breath-hold and resting-state acquisitions: (2.8 mm)^3^ isotropic voxels; 40% slice gap; 232 x 232 mm^2^ field of view; TE 105 ms; in-plane acceleration factor 2; partial Fourier factor 6/8; multiband factor 3. Breath-hold acquisitions comprised 21 slices with TR = 1000 ms, giving partial brain coverage which omitted the cerebellum and the inferior parts of the temporal lobes. To allow whole-cortex coverage, resting-state acquisitions comprised 24 slices with TR = 1100 ms. For each dfMRI acquisition, two b = 0 volumes were acquired, followed by alternating volumes with b = 200 and 1000 s mm^−2^ (Figure 1B). Two additional (b = 0) volumes were acquired with reverse phase encoding for correction of *B*_0_ field inhomogeneity distortions.

Multi-echo BOLD fMRI data (Kundu et al., 2017) were acquired with a gradient echo EPI sequence (TE 12.60, 30.18, 47.76, 65.34 ms) with flip angle 62°. All resolution and acceleration parameters were matched to the ADC-fMRI acquisitions. Two additional volumes were acquired with reverse phase encoding for correction of *B*_0_ field inhomogeneity distortions.

### 2.3 Physiological Monitoring

Expired CO_2_ pressure was recorded via a nasal cannula and ML206 gas analyser (ADInstruments, Sydney, Australia) via a BIOPAC MP160 physiological monitoring system (BIOPAC Systems Inc, Goleta, California, USA) which also recorded scanner triggers. AcqKnowledge data acquisition software (BIOPAC Systems Inc) was used to record signals sampled at 5 kHz, starting before and continuing after each scan to allow temporal shifting.

### 2.4 Data Analysis

#### 2.4.1 MRI Data

T_1_-weighted images were denoised using spatially adaptive non-local means filtering with ANTs (Manjón et al., 2010), then skull-stripped with Synthstrip (Hoopes et al., 2022) and segmented into tissue maps using FSL Fast (Zhang et al., 2001). Grey and white matter masks were transformed to the image space of each functional acquisition (ADC-fMRI and BOLD-fMRI) using rigid-body registration with ANTs in order to assess respiration-induced signals in grey and white matter. For group-level and between-subject analysis, T_1_-weighted images were registered to MNI standard space with nonlinear registration using ANTs.

To derive ADC-fMRI timecourses, dfMRI data were preprocessed according to Spencer et al. (2025), as follows. Magnitude image denoising was applied to the b = 200 s mm^−2^ and the b = 1000 s mm^−2^ timeseries using NORDIC (Moeller et al., 2021), with a 7x7x7 kernel and step size 1 for both g-factor estimation and PCA denoising. Gibbs unringing was applied to all volumes using MRtrix3 (Kellner et al., 2016; Tournier et al., 2019). The susceptibility off-resonance field was estimated from b = 0 volumes, and subsequently used to correct susceptibility-induced distortions in all volumes, using FSL Topup (Andersson et al., 2003; Jenkinson et al., 2012). Each b-value timeseries was corrected for motion using ANTs (Tustison et al., 2021), then registered to the initial b = 0 volume. A brain tissue mask was created from the Topup-corrected b = 0 volumes using Synthstrip (Hoopes et al., 2022) and used to remove non-brain voxels from the full timeseries. Subsequent analysis was performed on the ADC-fMRI timeseries calculated using Equation 1 in addition to each b-value timeseries (b200-dfMRI and b1000-dfMRI).

For BOLD-fMRI data, motion correction parameters were calculated on the timeseries of the first echo, then these transformations were used to correct the timeseries for all echoes, using ANTs. An optimally combined signal was calculated from a weighted average of echoes using Tedana, including denoising with TEDPCA and TEDICA (DuPre et al., 2021). Independent components from TEDICA denoising were visually inspected for each scan. For breath-hold data, the default TEDICA pipeline was found to remove BOLD components which were associated with the task (since hypercapnia induced large, widespread BOLD fluctuations). We therefore modified the default decision tree to include the task block design as a regressor, in order to keep any BOLD components which correlated with the breath-hold task. For resting-state data, the default TEDICA denoising decision tree was used, as this was found to only remove the expected non-BOLD, TE-independent noise components (likely because resting breathing fluctuations only result in small BOLD signal changes). For completeness, we demonstrated that the analysis with and without Tedana ICA denoising had minimal impact on the association with respiration (see Supplementary Materials -“BOLD Denoising”).

FSL Topup was applied to the first echo to calculate the susceptibility off-resonance field, then this was used to correct distortions in the optimally combined data. A brain mask was created using Synthstrip to remove non-brain voxels. For quality control of functional runs, absolute displacement from the first volume and framewise displacement between volumes (Power et al., 2012) were calculated from motion correction parameters for BOLD-fMRI, and for the b = 200 and b = 1000 s mm^−2^ timeseries from ADC-fMRI. Runs with a maximum absolute displacement greater than one voxel or a mean framewise displacement greater than 0.2 mm were rejected. In one subject, linear interpolation was used to replace a single outlying volume in the ADC-fMRI resting-state run. Two BOLD-fMRI resting-state runs and two ADC-fMRI resting-state runs were rejected due to excessive movement, in addition to one BOLD-fMRI breath-hold run.

All fMRI data (BOLD-fMRI, ADC-fMRI, b200-dfMRI, b1000-dfMRI) were spatially smoothed with a 4 mm FWHM Gaussian kernel, followed by high-pass temporal filtering at 0.01 Hz with a fourth order Butterworth filter. The preprocessing pipeline for all fMRI data is summarised in Supplementary Figure 1.

#### 2.4.2 p_**ET**_**CO**_**2**_

CO_2_ signals were low-pass filtered at 1 Hz with a fourth order Butterworth filter, then end-tidal peaks were identified with a peak detection algorithm and manually reviewed to remove erroneous measurements from partial exhales. Linear interpolation was performed between peaks to create the p_ET_CO_2_ timeseries, which were downsampled to 50 Hz to reduce file sizes. Each p_ET_CO_2_ timeseries was detrended with third order polynomials and convolved with the two-gamma variate canonical haemodynamic response function (Friston et al., 1998). All subsequent analysis was performed using the detrended, convolved p_ET_CO_2_ timeseries (Figure 1A).

For quality control, CO_2_ traces were visually inspected to identify unreliable measurements (for example, caused by the participant breathing through their mouth) or irregular breathing patterns. Breath-hold runs were additionally rejected if the participant failed to adequately comply with the task (for example, not completing the breath-hold period, or failing to exhale sufficiently to provide p_ET_CO_2_ at the beginning or end of the breath-hold period). Out of the runs remaining after rejecting those with excessive motion, p_ET_CO_2_ quality control led to the rejection of three (out of 13) ADC-fMRI resting-state runs, two (out of 13) BOLD-fMRI resting-state runs, ten (out of 30) ADC-fMRI breath-hold runs, and seven (out of 29) BOLD-fMRI breath-hold runs.

### 2.5 Noise Simulation

To investigate whether differences between ADC-fMRI and BOLD-fMRI in the association with p_ET_CO_2_ simply arise due to the lower signal-to-noise ratio (SNR) of ADC-fMRI (Spencer et al., 2025), we generated BOLD-fMRI timeseries with amplified noise levels. The raw BOLD-fMRI data from each breath-hold run was first denoised using NORDIC (Moeller et al., 2021) (with the parameters described for ADC-fMRI above) to generate a denoised timeseries and a residual timeseries for each echo time. The temporal order of the residuals was then randomly permuted in order to remove the temporal association between the task and any potential signal components contained in the residuals, whilst maintaining realistic noise statistics. These permuted residuals were then multiplied and added to the denoised data to provide a noise-amplified version of each echo time. The noised data was then preprocessed as described above for BOLD-fMRI, but without the Tedana denoising step when optimally combining echoes (in order to maintain the low SNR of the noise-amplified data), as shown in Supplementary Figure 1. This procedure was used to generate data with 4x, 8x and 16x the original noise levels. We calculated the image SNR of each dataset as the mean denoised signal divided by the standard deviation of the components removed by denoising (residuals). We also calculated the temporal SNR in the first baseline block (such that the signal is not impacted by any task-related fluctuations) as the temporal mean divided by the temporal standard deviation in the preprocessed data. In the 8x noised dataset, the image SNR of the first echo was lower than that of b200-dfMRI, and the image SNR of the second to fourth echoes were lower than that of b1000-dfMRI (Supplementary Figure 2 and Supplementary Table 1). In the 16x noised dataset, the SNR of all four echoes was lower than that of both b200-dfMRI and b1000-dfMRI. Both the 8x and 16x noised datasets had lower temporal SNR than ADC-fMRI (Supplementary Figure 3 and Supplementary Table 2).

### 2.6 Cross-correlation Analysis

We aimed to assess the relationship between fMRI contrasts and respiration, as characterised by p_ET_CO_2_, for each breath-hold and resting-state run. Due to between-subject differences in gas analyser delay and haemodynamic response (Murphy et al., 2011), in addition to within-subject variations in the haemodynamic response across the brain (Birn, Smith, et al., 2008; Chang & Glover, 2009; Zvolanek et al., 2023), we measured the cross-correlation in each voxel with time shifts of -30 s to0s at 0.5 s increments (this accounts for expected gas analyser delay of around 15 s). The peak correlation yielded the voxel-wise delay. Voxels with a peak correlation occurring at the extremes of the tested lags (-30 or 0) were excluded, as they cannot be deemed to be local maxima. To allow averaging across subjects, the peak correlation (*r*_*max*_) for each voxel was converted to a z-score using 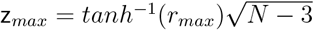, where N is the number of time points (Chang & Glover, 2009). The inclusion of 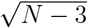 here adjusts the z-score for the different variance due to different numbers of time points included in the cross-correlation at different lags. To allow comparison between subjects, the latency map for each subject was shifted by the median latency of grey matter voxels. Maps of z_*max*_, *r*_*max*_ and latency were transformed to MNI space for group-level comparison.

### 2.7 Statistical Analysis

To summarise the association between fMRI contrast and p_ET_CO_2_, and to allow statistical comparison across subjects, we applied a threshold (z_*thresh*_) and measured number of voxels with a z-score *>*z_*thresh*_, and the variance explained (*r*_*max*_^2^) averaged across all voxels. For breath-hold, we first averaged the maps between available runs for a given subject before calculating the number of significant voxels and the variance explained. Pairwise comparisons of summary metrics were performed between contrasts using two-tailed unpaired Mann-Whitney U-tests with Bonferroni correction. For breath-hold analysis, we used z_*thresh*_ = 3.09, corresponding to p = 0.001. For resting-state analysis, z_*thresh*_ was determined for each contrast using surrogate analysis, described below. For completeness, we also analysed resting-state and breath-hold results with z_*thresh*_ values in the range 2.0–3.5.

We also measured the reproducibility of z_*max*_ maps within and between subjects. Similarity between two z_*max*_ maps was measured as the z-transformed Pearson correlation coefficient between voxel-wise values within the brain. In breath-hold data, we measured intra-subject similarity between the z_*max*_ maps of the two breath-hold runs. We then measured the inter-subject similarity by first averaging the z_*max*_ maps between available runs for a given subject, then measuring the mean similarity to every other subject. In resting-state data, we measured inter-subject similarity between z_*max*_ maps across subjects. We further assessed the within-subject similarity between the resting-state z_*max*_ map and the breath-hold z_*max*_ map.

#### 2.7.1 Surrogate Analysis

As cross-correlation analysis can inflate z-scores, and spurious fluctuations may impact cross-correlation values differently for each contrast, we derived contrast-specific significance thresholds from resting-state data to compare the summary statistics described above. Significance thresholds were derived from an empirical distribution of correlation values between surrogate p_ET_CO_2_ signals and the fMRI signal, under the null hypothesis that there is no correlation (Theiler et al., 1992). Similar to Chang and Glover (2009) and Yuan et al. (2013), we generated a set of 5000 surrogate signals based on the preprocessed p_ET_CO_2_ timeseries from one subject, and performed the cross-correlation between these 5000 surrogate signals and 5000 randomly selected voxels from each of the other subjects. Specifically, the surrogate timeseries were computed by applying a Fourier transform to the reference p_ET_CO_2_ timeseries, randomly shuffling the phases of the Fourier coefficients while keeping their magnitudes unchanged, and applying the inverse Fourier transform. We then repeated this step, taking in turn each subject p_ET_CO_2_ timeseries as the reference for the surrogate signal generation. Finally, we pooled the correlations to obtain the complete null distribution across all subjects. For each contrast, the significance threshold for a correlation between p_ET_CO_2_ and fMRI timeseries was calculated as the z_*max*_ value corresponding to p *<* 0.05 (one tail) under this null distribution. The resulting significance thresholds were: 4.42 (BOLD-fMRI); 2.84 (b200-dfMRI); 2.36 (b1000-dfMRI); 2.48 (ADC-fMRI) (Supplementary Figure 4). Note that this analysis was not performed on the breath-hold data, as the p_ET_CO_2_ timeseries for every subject is dominated by the task frequency, therefore the surrogate signals would still have a high correlation with the original timeseries. Instead, for breath-hold analysis, we used a fixed z_*thresh*_ = 3.09, corresponding to p = 0.001 for all functional contrasts, as described in *Statistical Analysis*.

## 3 Results

### 3.1 Breath-Hold Response

The association between p_ET_CO_2_ and each functional contrast was measured during a breath-hold experiment. As shown in the group-average z_*max*_ map, BOLD-fMRI displays a strong association with p_ET_CO_2_, mostly in grey matter, but also with less intensity in white matter (Figure 2A). A similar but less intense pattern is also observed in grey matter in b200-dfMRI and b1000-dfMRI, while most white matter voxels have a z-score below 1.5. For ADC-fMRI, the voxels showing an association between p_ET_CO_2_ and functional signals are scattered throughout the brain, with some possible patterns overlapping with the ventricles and the cortical ribbon in close proximity to the cerebrospinal fluid (CSF). To investigate the spatial variations in the vascular response, the time lags corresponding to each z_*max*_ are plotted as group-average latency maps (Figure 2B). Subject-level z_*max*_ and latency maps are shown in Supplementary Figure 5A, and the median latency used to normalise each subject is shown in Supplementary Figure 6. The BOLD-fMRI latency map shows a clear pattern of longer delays in white matter compared to grey matter (Figure 2B), indicating a delayed haemodynamic response in white matter. Conversely, the b200-dfMRI, b1000-dfMRI and ADC-fMRI latency maps show no discernible pattern.

**Figure 2.**
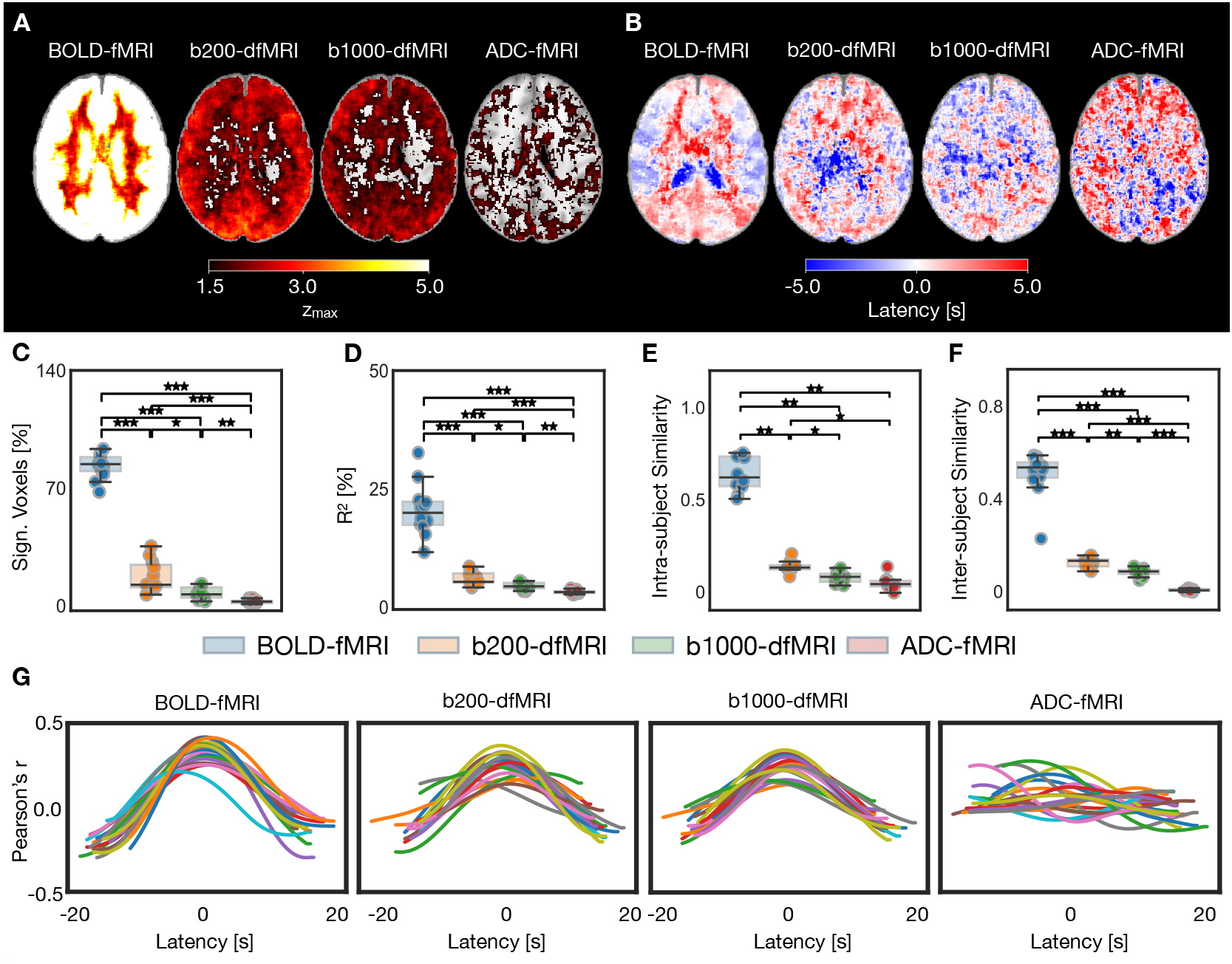
**Breath-hold: p**_**ET**_**CO**_2_ **vs fMRI association**, for BOLD-fMRI, b200-dfMRI, b1000-dfMRI and ADC-fMRI. BOLD-fMRI (n = 12, 22 runs), b200-dfMRI, b1000-dfMRI and ADC-fMRI (n = 11, 20 runs). A) Group-average z_*max*_ maps. B) Group-average latency maps. C) Significant voxels (z_*max*_ *>* 3.09) as a percentage of voxels in the brain. D) Coefficient of determination (percentage variance explained) in all voxels. E) Intra-subject similarity in breath-hold z_*max*_ maps between task runs. F) Intersubject similarity in breath-hold z_*max*_ maps; each point represents the mean similarity from the z_*max*_ map of one subject (averaged across available runs) to all other subjects. G) Correlation profiles: each line corresponds to one breath-hold run, showing the mean Pearson’s correlation coefficient (averaged across voxels with a significant z_*max*_) over the range of lags tested, shifted to the median grey matter latency. Brain maps show MNI slice z = 95. Two-sided Mann-Whitney U-tests with Bonferroni correction were performed. *p *<* 0.05; **p *<* 0.01; ***p *<* 0.001.

The association between p_ET_CO_2_ and each functional contrast is further described statistically with the percentage of significant voxels (Figure 2C), using a z-threshold of 3.09 (p *<* 0.001), and the percentage variance explained (Figure 2D). The percentage of significant voxels across the brain decreases significantly from one contrast to the next (BOLD-fMRI *>* b200-dfMRI *>* b1000-dfMRI *>* ADC-fMRI, Figure 2C). The percentage of significant voxels with incremental z-thresholds ranging from 2.0 to 3.5 is shown in Supplementary Figure 7A.

To determine the extent to which the association between p_ET_CO_2_ and each functional contrast reflects an underlying dependence on vascular signals rather than spurious correlations, we measured the intrasubject similarity between z_*max*_ maps from pairs of breath-hold runs in the same subject (Figure 2E), and inter-subject similarity between z_*max*_ maps from different subjects (Figure 2F). Intra- and inter-subject similarity was low (*<* 0.3) for b200-dfMRI, b1000-dfMRI and ADC-fMRI. ADC-fMRI had significantly lower intra-subject similarity than b200-dfMRI and BOLD-fMRI (p *<* 0.05), and than b1000-dfMRI (p *<* 0.07, non-significant trend). Inter-subject similarity was significantly different (p *<* 0.05) between all contrasts (BOLD-fMRI *>* b200-dfMRI *>* b1000-dfMRI *>* ADC-fMRI).

Finally, to visualise whether the peak cross-correlations were temporally distinct, subject-level correlation profiles are displayed in Figure 2G, showing the average correlation coefficient over significant voxels for the full range of latencies tested. While BOLD-fMRI, b200-dfMRI and b1000-dfMRI show distinct peaks in the correlation profiles, the ADC-fMRI correlation profiles are not synchronised, indicating a lack of coherence across the brain in the association with p_ET_CO_2_. This is consistent with expected residual T_2_ contributions to b200 and b1000 timecourses, that are minimal in the ADC timecourse.

To ensure that the BOLD-fMRI z-scores were not artificially inflated because the BOLD-fMRI timeseries contained twice as many time points as the diffusion timeseries, we repeated the analysis after downsampling the BOLD-fMRI timeseries by keeping every other time point. Similar results were obtained (Supplementary Figure 8A-D), supporting that the stronger association between respiration and BOLD-fMRI compared to the diffusion functional contrasts was not due to the sampling rate.

### 3.2 Noise-Amplified Data

To assess whether the lower association between p_ET_CO_2_ and b200-dfMRI, b1000-dfMRI and ADC-fMRI was simply due to the reduced SNR of the dfMRI sequence, we created noise-amplified versions of all BOLD-fMRI breath-hold datasets with 4x, 8x and 16x the original noise levels. Increasing the noise levels decreased the group-average z-score (Supplementary Figure 9A), the percentage of significant voxels (Supplementary Figure 9B) and the variance explained (Supplementary Figure 9C). However, the association with respiration remained much stronger than that of ADC-fMRI (Figure 9A). Despite the reduced z-scores in noised data, the intra-subject similarity (Supplementary Figure 9D) and inter-subject similarity (Supplementary Figure 9E) remained high.

### 3.3 Resting Breathing Response

As breath-holding is an extreme case of CO_2_ variation, we further aimed to determine how spontaneous fluctuations in respiratory rate and depth occurring during resting breathing could affect the functional contrasts (Figure 3). A similar pattern is seen in the BOLD-fMRI z_*max*_ map (Figure 3A), while the p_ET_CO_2_ signal is less uniformly associated with the b200-dfMRI and b1000-dfMRI signals in grey matter. In white matter, almost no voxels show a z_*max*_ greater than 1.5 in b200-dfMRI and b1000-dfMRI. This holds across the whole brain in ADC-fMRI. Similar to breath-hold, BOLD-fMRI resting breathing latency maps show a separation between grey and white matter latencies, with longer latencies in the latter (Figure 3B). This pattern is visible in six out of eleven subjects in BOLD-fMRI (Supplementary Figure 5B) and is also partially discernible in b200-dfMRI.

**Figure 3.**
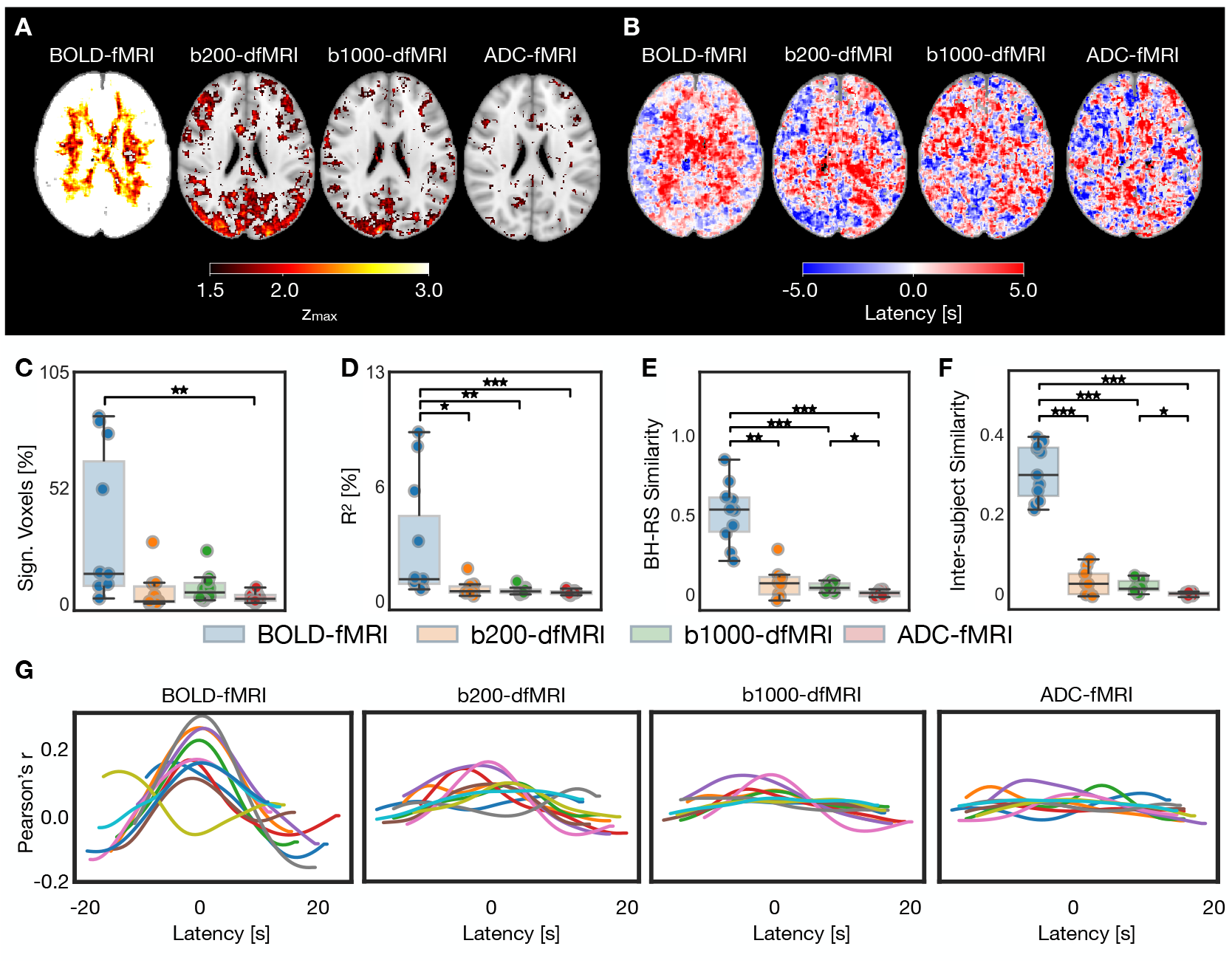
**Resting-state: p**_**ET**_**CO**_2_ **vs fMRI association**, for BOLD-fMRI, b200-dfMRI, b1000-dfMRI and ADC-fMRI. BOLD-fMRI (n = 11), b200-dfMRI, b1000-dfMRI and ADC-fMRI (n = 10). A) Group-average z_*max*_ maps. B) Group-average latency maps. C) Significant voxels as a percentage of brain voxels. D) Coefficient of determination (percentage variance explained) in all voxels. E) Reproducibility of z_*max*_ maps between breath-hold and resting-state experiments, measured by within-subject similarity between breath-hold z_*max*_ maps (averaged across available runs) and resting-state z_*max*_ maps. F) Intersubject similarity in resting-state z_*max*_ maps; each point represents the mean similarity from the z_*max*_ map of one subject to all other subjects. G) Correlation profiles: each line corresponds to one subject, showing the mean Pearson’s correlation coefficient (averaged across voxels with a significant z_*max*_) over the range of lags tested, shifted to the median grey matter latency. The following contrast-specific z-thresholds, derived from surrogate analysis, were used to determine voxel significance: 4.42 (BOLD-fMRI); 2.84 (b200-dfMRI); 2.36 (b1000-dfMRI); 2.48 (ADC-fMRI). Brain maps show MNI slice z = 95. Pairwise two-sided Mann-Whitney U-tests with Bonferroni correction were performed. *p *<* 0.05; **p *<* 0.01; ***p *<* 0.001.

As expected from the more subtle variations in CO_2_ observed at rest, fewer voxels show a significant association between p_ET_CO_2_ and functional contrast (Figure 3C). However, BOLD-fMRI still shows a significantly larger percentage of significant voxels than ADC-fMRI, and a larger variance explained than the dfMRI and ADC-fMRI contrasts (p *<* 0.05, Figure 3D). The statistical significance thresholds used in Figures 3C were determined using surrogate analysis, but the same analysis was performed with incremental z-thresholds ranging from 2.0 to 3.5 (Supplementary Figure 7B).

To quantify the extent to which the resting-state data captured the same vascular response as the breath-hold experiments, we measured the within-subject similarity between breath-hold z_*max*_ maps and resting-state z_*max*_ maps (Figure 3E), and the inter-subject similarity between resting-state z_*max*_ maps from different subjects (Figure 3F). Similarity with breath-hold z_*max*_ maps, and similarity between subjects, was higher for BOLD-fMRI than for dfMRI and ADC-fMRI (p *<* 0.05). ADC-fMRI was lower than b1000-dfMRI (p *<* 0.05) and than b200-dfMRI (non-significant trends: p *<* 0.2 for similarity with breath-hold; p *<* 0.09 for inter-subject similarity).

Finally, the subject-level correlation profiles also show coherent profiles, albeit with lower amplitude, for BOLD-fMRI (Figure 3G). The correlation profiles show less coherence for b200-dfMRI and b1000-dfMRI, and are flat for ADC-fMRI.

Similar to the breath-hold analysis, we performed the same analysis on the downsampled BOLD timeseries (Supplementary Figure 8E-H). Similar conclusions can be drawn for the downsampled BOLD-fMRI.

## 4 Discussion

In this study, we aimed to address concerns that the ADC-fMRI contrast may be driven by neurovascular coupling, by evaluating its association with non-neuronal haemodynamic fluctuations. We used a breathhold task to induce hypercapnia, which in turn causes an increase in the BOLD signal due to the vasodilatory effects of CO_2_. The minimal correlation between p_ET_CO_2_ and ADC-fMRI timecourses in this experiment, and the low intra- and inter-subject reproducibility in z_*max*_, suggest the ADC-fMRI contrast is independent from vascular contributions. Therefore, the T_2_-weighting of the dfMRI signal (and its sensitivity to vascular contributions) is suppressed when taking the ratio between two b-values in the ADC-fMRI acquisition.

The association between respiration and BOLD-fMRI is confirmed by strong correlations with p_ET_CO_2_. Further, the high intra- and inter-subject reproducibility in z_*max*_ from BOLD-fMRI suggests the same underlying physiological mechanisms for the association between respiration and BOLD signal are captured within and between subjects. Both b200-dfMRI and b1000-dfMRI remain sensitive to vascular contributions due to T_2_-weighting, as highlighted by the previously reported similarities between BOLD and dfMRI spatial maps during visual and motor tasks, as well as a slow return to baseline observed in Spencer et al. (2025). However, this vascular association is weaker than BOLD-fMRI, as evidenced by weaker correlations with p_ET_CO_2_, and lower similarity metrics suggesting the spatial patterns of z_*max*_ from these contrasts are less rooted in an underlying association between physiology and contrast. This reduced association with respiration is likely due to the substantial suppression of direct contributions from the blood water to the signal, using b-values ≥ 200 s mm^−2^. In particular, diffusion weighting effectively suppresses intravascular contributions (which accounts for ∼50% of the SE signal at 3T, Poser and Norris 2007) while providing sensitivity to microstructural fluctuations (De Luca et al., 2019; Le Bihan, 2012; Le Bihan et al., 2006), as evidenced by an earlier diffusion response onset than the BOLD response (Aso et al., 2009, 2013; Nguyen-Duc et al., 2025; Nunes et al., 2021; Spencer et al., 2025; Williams et al., 2014, 2015). Consistent with this suppression of intravascular contributions, the patterns of association between p_ET_CO_2_ and b200-dfMRI are less pronounced with b1000-dfMRI, but remain detectable. In addition to direct blood signal suppression, the association with both b200-dfMRI and b1000-dfMRI is likely also lower as compared to our BOLD-fMRI timeseries due to the lower sensitivity to magnetic susceptibility changes of the spin-echo (T_2_-weighting) vs the gradient-echo (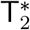-weighting) sequence. ADC-fMRI showed minimal association with vascular signals with low reproducibility in correlation maps. This suggests further suppression of T_2_ BOLD contrast in the ADC estimation from two subsequent diffusion-weighted images, and the near absence of vascular sources in the ADC-fMRI time courses in brain voxels. For ADC-fMRI, there was large variance in the latency values across grey matter voxels in each subject (Supplementary Figure 6C and D), suggesting the correlations are mostly spurious with arbitrary latencies. Conversely, the strong correlation between BOLD-fMRI and p_ET_CO_2_ allows more precise measurement of the latency in most grey matter voxels (reflected by the low variance in latency values within each subject). The resulting median latency values for BOLD-fMRI capture the variability between subjects due to physiology, breathing depth, or experimental setup.

In the ADC-fMRI spatial maps, there appeared to be some weak correlation patterns in the ventricles and the cortical ribbon close to the CSF. A possible mechanism underlying this association with CSF is the “Monro–Kellie hypothesis” (Greitz et al., 1992), whereby CSF volume decreases to compensate for the increased blood flow and volume following the haemodynamic response (Piechnik et al., 2009). This has been reported to induce changes in the CSF volume fraction of -0.6% in the visual cortex following visual stimulation (Donahue et al., 2006; Jin & Kim, 2010). According to the three compartment model (Rydhög et al., 2017), a -0.6% change in CSF volume fraction would result in a measured ADC change of at most -0.3% and likely much less (see Supplementary Materials – “CSF Contribution”). In addition to the change in CSF volume fraction, a decrease in CSF mobility from compression may decrease the ADC of the CSF compartment (and therefore further decrease the overall ADC). However, we estimate the measured ADC change resulting from this effect to be smaller than -0.02% (see Supplementary Materials – “CSF Contribution”). Consistent with our estimations, we found no evidence of a negative association between ADC and p_ET_CO_2_ (Figures 2G and 3G, and Supplementary Figure 11). Spencer et al. (2025) reported an ADC-fMRI signal decrease of around 1% during a visual task, which could not be solely explained by the CSF compensation mechanisms above.

BOLD-fMRI datasets with noise amplified to levels comparable to the ADC-fMRI acquisitions maintained strong association with respiration and high reproducibility of correlation maps, demonstrating that the lower dependence of ADC-fMRI on p_ET_CO_2_ is not simply reflective of the lower SNR. The reduced dependence of dfMRI and ADC-fMRI on p_ET_CO_2_ was still evident when the subjects were breathing freely in a resting-state experimental setting more representative of conventional fMRI study conditions.

In the latency maps from both breath-hold and resting-state, a clear separation between grey and white matter can be seen for BOLD-fMRI, with the response to respiration occurring later in white matter. In resting-state, this pattern is also visible to a lesser extent in the subject-level maps for b200-dfMRI and b1000-dfMRI, with some subjects exhibiting higher z-scores or earlier lags preferentially located in the grey matter. A delayed association between p_ET_CO_2_ and BOLD-fMRI in white matter was previously been reported in breath-hold tasks (Champagne et al., 2019; Thomas et al., 2014; Zvolanek et al., 2023) and in resting-state (Tong et al., 2017), which can be explained by longer (up to 3 s, Liu et al. 2012) transit time to reach white matter due to vascular geometry (Özbay et al., 2018; Thomas et al., 2014), poorer vascular reactivity in white matter (Champagne et al., 2019), or longer time for extravascular CO_2_ to build up due to lower cerebral blood flow (Thomas et al., 2014). In addition to the large difference in latency between grey and white matter, the latency varies by several seconds across the cortex, demonstrating the heterogeneity of the haemodynamic response (Handwerker et al., 2004; Li et al., 2019). This challenges the use of the canonical haemodynamic response function – which assumes a uniform BOLD response across the brain – as well as traditional resting-state functional connectivity analysis – which assumes that co-varying neural activity is captured by instantaneous correlations in the BOLD response of distant brain regions. Conversely, ADC-fMRI relies on neuromorphological coupling and may provide increased temporal synchronicity, which overcomes the inherent biases of BOLD-fMRI related to the heterogeneity in the haemodynamic response.

Furthermore, the influence of physiological signals on functional contrasts is a confound for the analysis of brain function. For example, BOLD-fMRI functional connectivity analysis of resting-state networks (RSNs) is particularly susceptible to bias from physiological noise. RSNs are assumed to arise from correlations in neuronal activity (Biswal et al., 1995; Greicius et al., 2003), as supported by electrical or magnetic recordings (Brookes et al., 2011; Laufs et al., 2003; K. J. Miller et al., 2009). However, the vascular response to spontaneous respiration fluctuations can induce “physiological connectivity” (Chen et al., 2020) – correlations in the BOLD signal across different brain regions, which may be mistaken for functional connectivity (Chen et al., 2020; Tong et al., 2015). The interlinked development of the vascular structure and neuronal network structure may lead to distant brain regions sharing similar vascular anatomy and reactivity to vasodilatory signals to efficiently support the increased metabolic demands of the network (Bright et al., 2020; Chen et al., 2020; Quaegebeur et al., 2011). While these distinct neurovascular signatures shared among remote brain regions may obscure the ability of BOLD-fMRI and dfMRI to detect co-varying neuronal activity, ADC-fMRI is robust to vascular signals and therefore has the potential to offer more unbiased investigation of RSNs (de Riedmatten et al., 2025).

A complete analysis of RSNs revealed by our ADC-fMRI data will be the focus of a separate, dedicated work. We underline however that, while ADC-fMRI timecourses are robust to vascular contamination, they do contain meaningful functional information. To this end, we briefly present supplementary results showing resting-state functional connectivity captured by the ADC-fMRI timecourse (see Supplementary Materials – “Resting-State Functional Connectivity”), providing evidence that ADC-fMRI is sensitive to resting-state activity. This is consistent with previous work by Spencer et al. (2025) using a similar contrast design, which showed that isotropic ADC-fMRI was able to detect neuronal activity during both visual and motor tasks. Similarly, ADC-fMRI with linear diffusion encoding was previously also shown to detect neuronal activity during visual (Nguyen-Duc et al., 2025; Nicolas et al., 2017; Spees et al., 2013) and motor tasks (Nguyen-Duc et al., 2025), and during resting-state (de Riedmatten et al., 2025).

The ADC-fMRI design in the present study builds on the acquisition used by Spencer et al. (2025) with the addition of cross-terms compensated waveform design (Szczepankiewicz & Sjölund, 2021) for a comprehensive mitigation of all three vascular contribution sources. Since this comes at the expense of a longer echo time and therefore a penalty in SNR, we compensated for this effect by acquiring larger voxels, and confirmed that the SNR is comparable between the two methods (Supplementary Figure 10 and Supplementary Table 3).

While ADC-fMRI in its current form does not match the temporal resolution and SNR of BOLD-fMRI, our results demonstrate that the underlying functional information is distinct from neurovascular coupling. Thus, with developments to hardware, acquisition or post-processing denoising to improve sensitivity and temporal resolution, it has the potential to overcome the intrinsic limitations of the BOLD contrast, providing a contrast that is more specific to neuronal signals.

### 4.1 Limitations

The breath-hold task is designed to elicit a vascular response independently of neuronal signals (Murphy et al., 2011; Pinto et al., 2021). In reality, there may be neuronal activation associated with the visual and cognitive processing of the task instructions (Birn, Smith, et al., 2008; Pinto et al., 2021). Additionally, breath-holding may modulate neuronal activity (as reported by studies using non-vascular neuroimaging approaches such as MEG (Driver et al., 2016; Kluger & Gross, 2021) and EEG (Yuan et al., 2013)). However, our results suggest limited modulation of neuronal activity during hypercapnia. The crosscorrelation analysis revealed an association between p_ET_CO_2_ and BOLD-fMRI at a delay encompassing the time taken for breath-holding to induce an accumulation of CO_2_ in the brain (Birn, Smith, et al., 2008; Murphy et al., 2011). While neuronal activation associated with performing the task may be temporally synchronised with the breath-hold task blocks, any influence of hypercapnia on neuronal activity would be synchronised with cerebral CO_2_ levels captured by the delayed p_ET_CO_2_ trace, and therefore would be inseparable from the purely vascular changes of non-neuronal origin. The correlation profiles for the breath-hold task demonstrate distinct peaks that were synchronised across subjects for BOLD-fMRI, b200-dfMRI and b1000-dfMRI, while ADC-fMRI showed flat correlation profiles with no coherence.

In this work, the hypercapnia challenge was implemented as a breath-holding task rather than a gas challenge, preventing direct control of the inhaled CO_2_ concentration. However, CO_2_ fluctuations were tracked using a gas analyser, allowing us to include p_ET_CO_2_ signals in the experimental design to control for inter-subject differences in task compliance and CO_2_ concentration.

## 5 Conclusions

This study provides evidence that ADC-fMRI, when carefully designed to mitigate cross-terms and perfusion effects, is largely free of vascular contamination, both in breath-holding task and under resting conditions. We conclude that previously reported ADC-fMRI results arise from sensitivity to microstructure fluctuations. Its diffusion timeseries counterparts, b200-dfMRI and b1000-dfMRI may offer a favourable balance between sensitivity to microstructural changes with good temporal resolution, and some vascular contribution. However, the residual structured physiological noise in these contrasts may still obscure true neuronal signals, reinforcing ADC-fMRI as the more optimal contrast for specifically capturing neuronal activity with minimal vascular confounds.

## Data and Code Availability

The data and associated code will be available upon publication.

## Author Contributions

Conceptualization: I.J., I.d.R., A.S.; Methodology: I.J., I.d.R., A.S.; Validation: I.d.R., A.S.; Formal analysis: I.d.R., A.S.; Investigation: A.S., J-B.P., J.N-D., I.d.R., O.E., F.S.; Data curation: I.d.R., A.S.; Writing-original draft: I.d.R., A.S.; Writing - Review & Editing: I.d.R, A.S., J-B.P., J.N-D., F.S., I.J., O.E.; Visualisation: I.d.R, A.S.; Supervision: I.J.; Funding acquisition: I.J.

## Acknowledgements

The authors thank Jean-Baptiste Ledoux for his help in setting up the fMRI experiments, Stefano Moia for advice regarding physiological monitoring, and Eneko Uruñuela for input on multi-echo denoising. They also thank Antoine Delattre-Klauser, Tobias Kober, Tom Hilbert, Gian Franco Piredda, and Xavier Sieber for their help with the dfMRI sequence. The authors acknowledge the CIBM Center for Biomedical Imaging for providing expertise and resources to conduct this study.

