## Supplementary Materials for "Evaluating the dependence of ADC-fMRI on haemodynamics in breath-hold and resting-state conditions"

Supplementary Table 1: **Average image SNR for raw and noise-amplified BOLD-fMRI and ADC-fMRI breath-hold data.** Mean (standard deviation) across subjects of the average SNR within the brain are shown for the four echo times (TE) from BOLD-fMRI data, and for b = 200 and b = 1000 s $mm^{-2}$ from the ADC-fMRI acquisition for comparison.

| | $TE_1$ | $TE_2$ | $TE_3$ | $TE_4$ |
|---|---|---|---|---|
| Raw BOLD-fMRI | 149.40 (11.98) | 111.55 (8.25) | 80.65 (6.11) | 57.89 (4.33) |
| 4x Noise BOLD-fMRI | 37.35 (3.00) | 27.89 (2.06) | 20.16 (1.53) | 14.47 (1.08) |
| 8x Noise BOLD-fMRI | 18.67 (1.50) | 13.94 (1.03) | 10.08 (0.76) | 7.24 (0.54) |
| 16x Noise BOLD-fMRI | 9.34 (0.75) | 6.97 (0.52) | 5.04 (0.38) | 3.62 (0.27) |
| b = | 200 | | 1000 | |
| ADC-fMRI | 31.66 (2.34) | | 15.44 (0.93) | |

Supplementary Table 2: **Average temporal SNR.** Mean (standard deviation) across subjects of the average temporal SNR within the brain are shown for BOLD-fMRI data at each amplified noise level, and for b200-dfMRI, b1000-dfMRI and ADC-fMRI.

| | Temporal SNR |
|---|---|
| Raw BOLD-fMRI | 94.47 (6.39) |
| 4x Noise BOLD-fMRI | 38.72 (2.50) |
| 8x Noise BOLD-fMRI | 21.06 (1.36) |
| 16x Noise BOLD-fMRI | 11.32 (0.81) |
| b200-dfMRI | 57.20 (4.53) |
| b1000-dfMRI | 37.82 (3.09) |
| ADC-fMRI | 22.96 (1.88) |

Supplementary Table 3: **Image SNR value comparison with previous data.** Mean (standard deviation) across subjects in the average image SNR within the brain are shown for the breath-hold data used in this study and visual task data from Spencer et al. (2025). The p-value of two-tailed t-tests is shown. *$p < 0.05$.

| b [s $mm^{-2}$] | Region | Breath-hold data | Visual task data | p |
|---|---|---|---|---|
| 200 | Whole brain | 28.75 (1.79) | 29.61 (1.76) | 0.1960 |
| | Cortex | 41.32 (2.59) | 41.58 (3.26) | 0.7999 |
| | Midbrain | 18.56 (1.35) | 19.41 (1.61) | 0.1184 |
| 1000 | Whole brain | 14.11 (0.66) | 14.85 (0.90) | 0.0115* |
| | Cortex | 20.04 (1.08) | 20.48 (1.35) | 0.3219 |
| | Midbrain | 8.93 (0.47) | 9.72 (0.88) | 0.0026* |

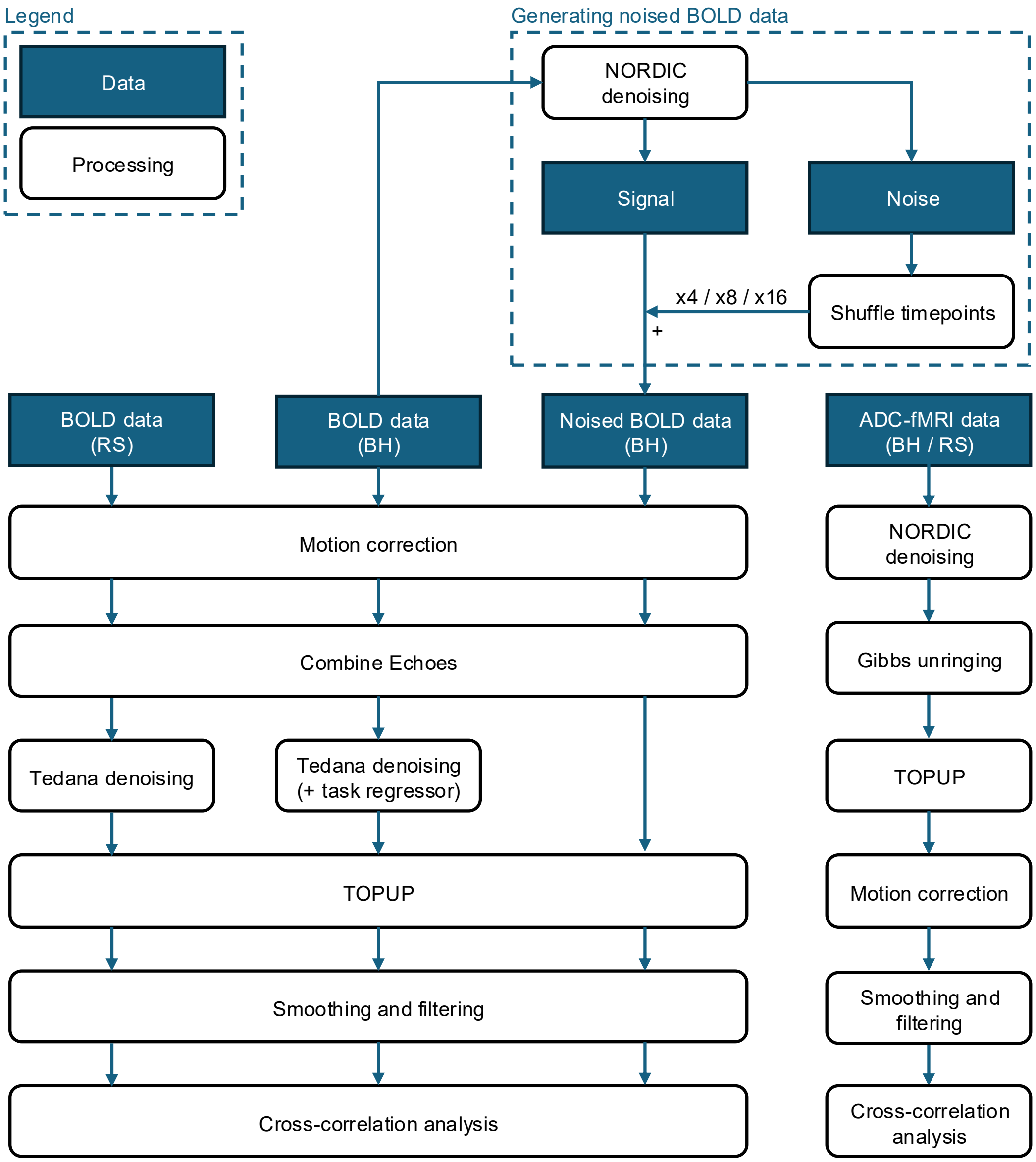


Supplementary Figure 1: **Preprocessing flowchart.** Preprocessing pipelines are shown for breath-hold (BH) and resting-state (RS) data, for BOLD-fMRI and ADC-fMRI (which comprises b200-dfMRI and b1000-dfMRI data).

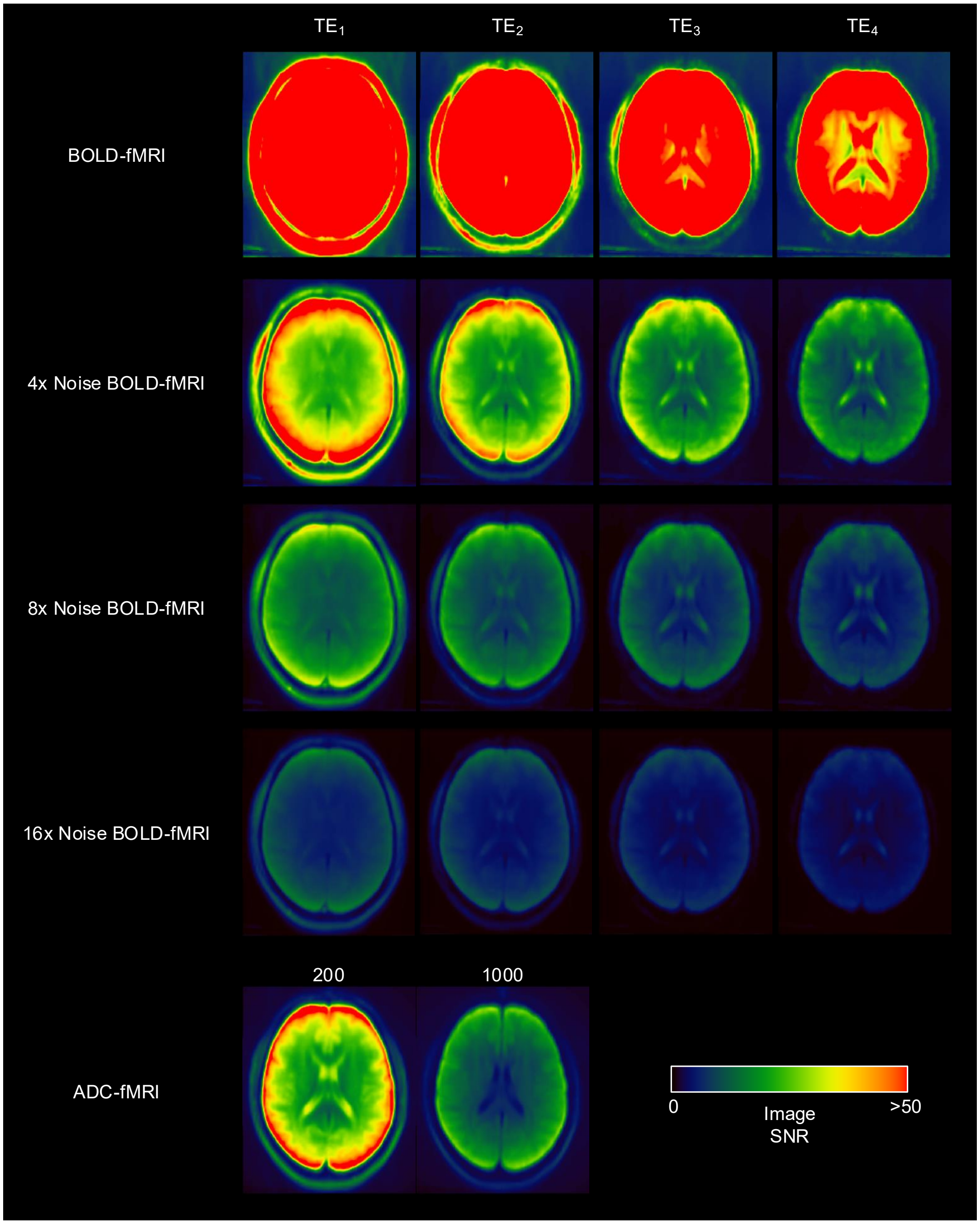


Supplementary Figure 2: **Group-mean image SNR maps for raw and noise-amplified BOLD-fMRI data, and ADC-fMRI data.** SNR was calculated as the mean denoised signal divided by the standard deviation of the components removed by denoising (residuals). To generate noised data, residuals were temporally randomly permuted and multiplied (4x, 8x or 16x) before adding them to the denoised data. Group-mean SNR maps are shown for the four echo times (TE) from BOLD-fMRI data, and for b = 200 and b = 1000 s mm$^{-2}$ from the ADC-fMRI acquisition for comparison. Mean SNR values within the brain are shown in Supplementary Table 1. Note that image SNR values cannot be calculated for ADC-fMRI, as this is derived from the constituent b-value acquisitions.

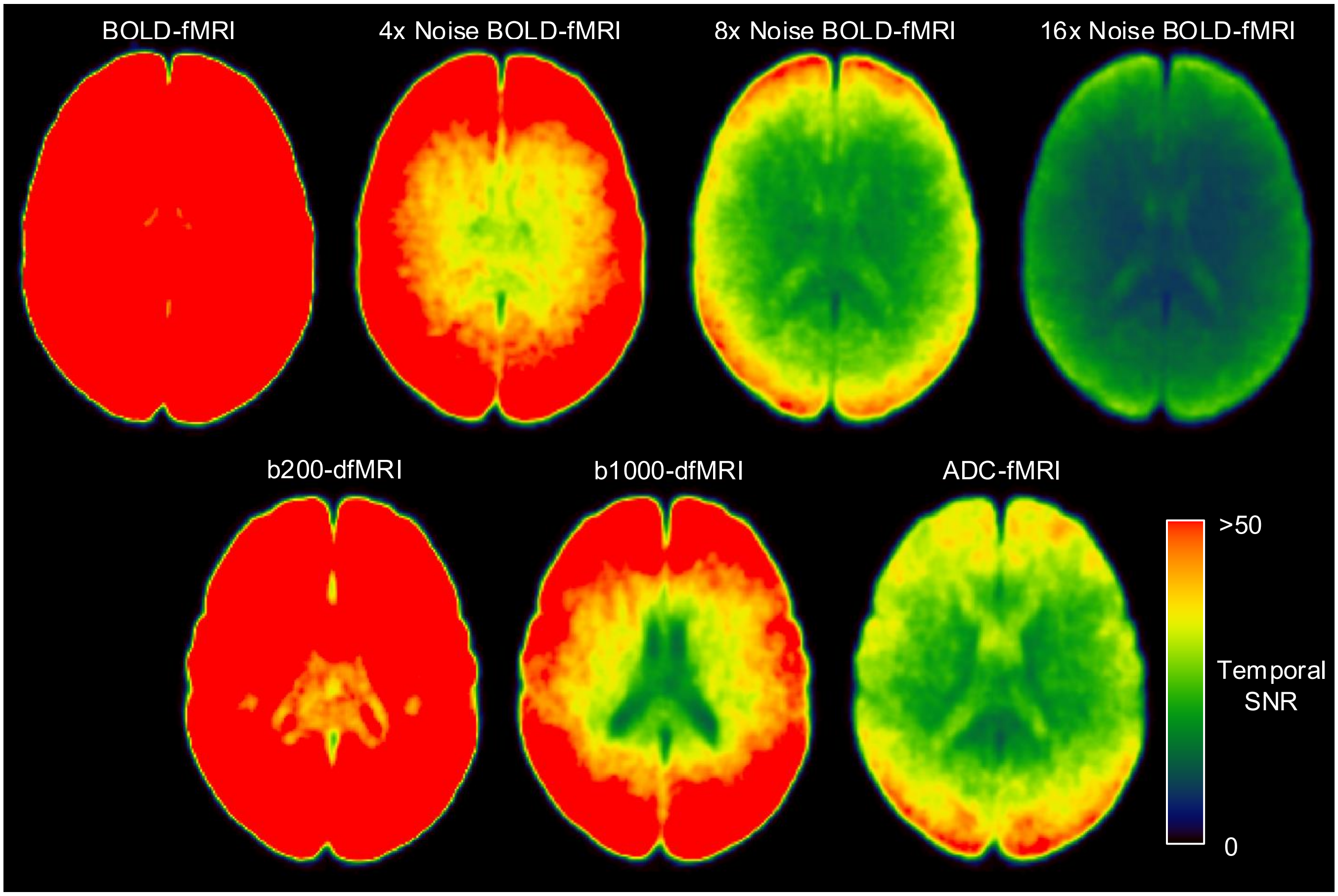


Supplementary Figure 3: **Group-mean temporal SNR maps for breath-hold data.** Temporal SNR was calculated in each voxel as the mean of the time course divided by the standard deviation. This was calculated from the first 18 s baseline block to remove any influence of task-associated signal fluctuations. BOLD-fMRI timeseries were downsampled so that the number of timepoints matched dfMRI and ADC-fMRI. Mean temporal SNR values within the brain are shown in Supplementary Table 2.

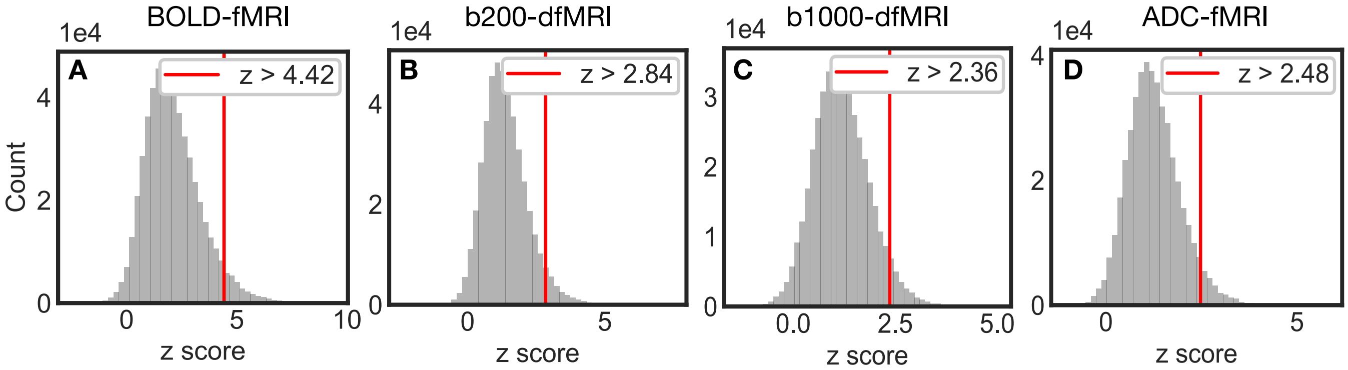


Supplementary Figure 4: **Significance threshold determination for resting-state experiment, using surrogate analysis** ($p < 0.05$, one tail), for BOLD-fMRI, b200-dfMRI, b1000-dfMRI and ADC-fMRI contrast. The histograms correspond to the generated null distribution, under the null hypothesis that the correlations between $p_{ET}CO_2$ and the fMRI signal are not significant.

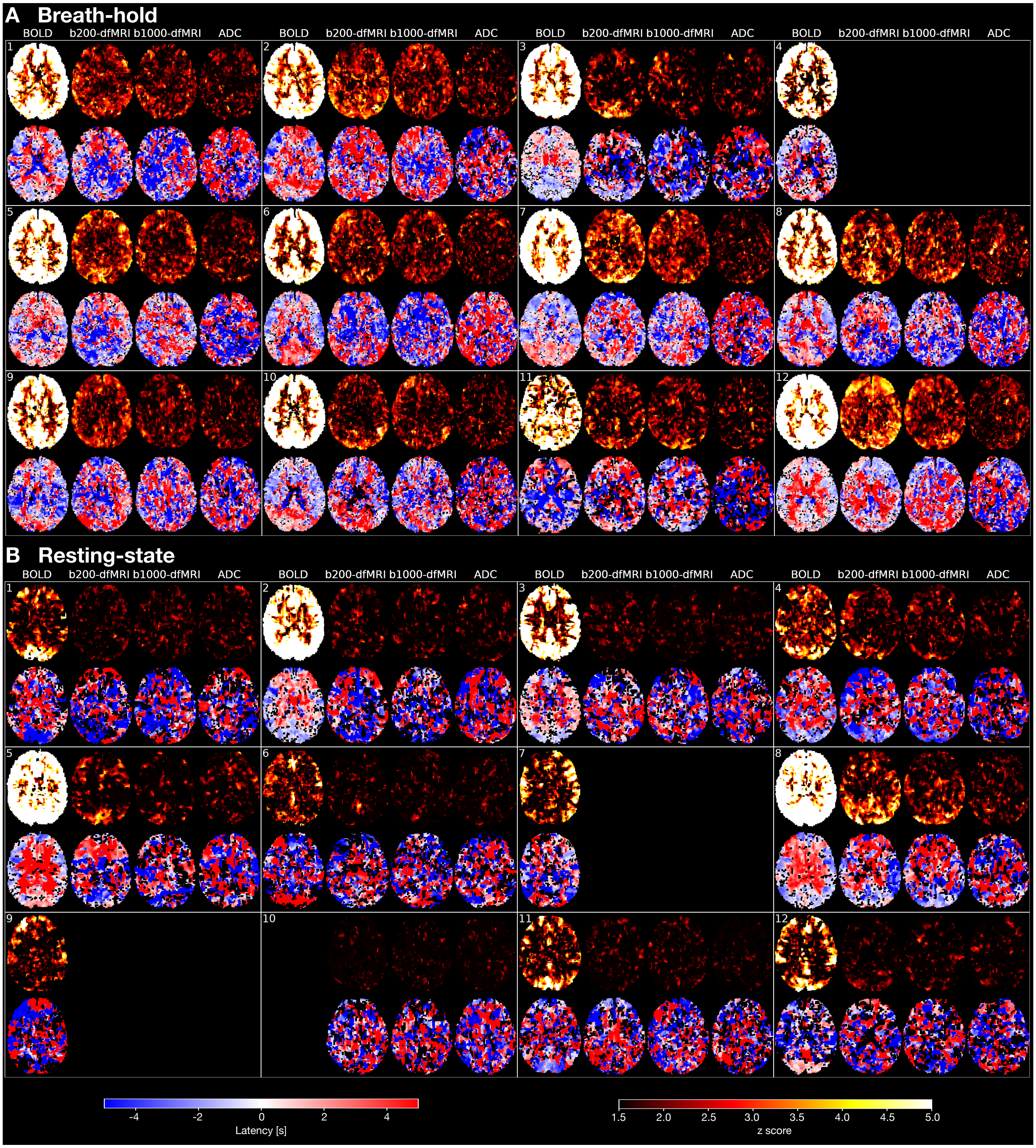


Supplementary Figure 5: **Individual z$_{max}$ and latency maps**. A) Breath-hold: BOLD-fMRI (n = 12, 22 runs), b200-dfMRI, b1000-dfMRI and ADC-fMRI (n = 11, 20 runs). B) Resting-state: BOLD-fMRI (n = 11), b200-dfMRI, b1000-dfMRI and ADC-fMRI (n = 10). Subjects are delimited by white rectangles. In each box, the first row corresponds to the z$_{max}$ maps, and the second row to the latency maps. Empty spaces correspond to missing data. Note that the subject numbering does not correspond between breath-hold and resting-state.

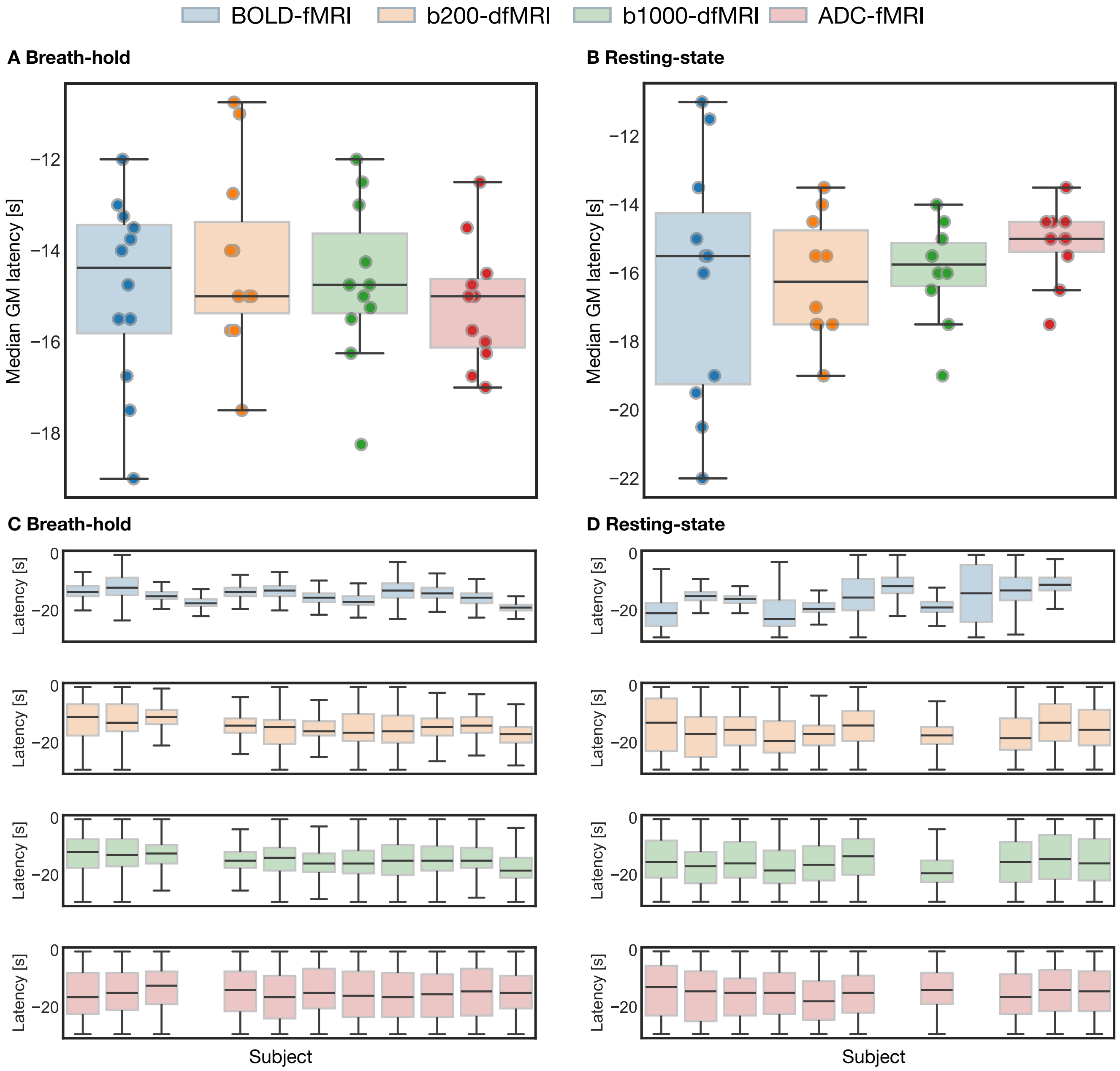


Supplementary Figure 6: **Grey matter latency values**. The median value per subject is plotted for the group: A) Breath-hold: BOLD-fMRI (n = 12, 22 runs), b200-dfMRI, b1000-dfMRI and ADC-fMRI (n = 11, 20 runs). B) Resting-state: BOLD-fMRI (n = 11), b200-dfMRI, b1000-dfMRI and ADC-fMRI (n = 10). For both the breath-hold task data and resting-state data, pairwise Mann-Whitney U-tests revealed no significant differences in median grey matter latency. The distribution of latency values across all grey matter voxels are also plotted for each subject: C) Breath-hold. D) Resting-state.

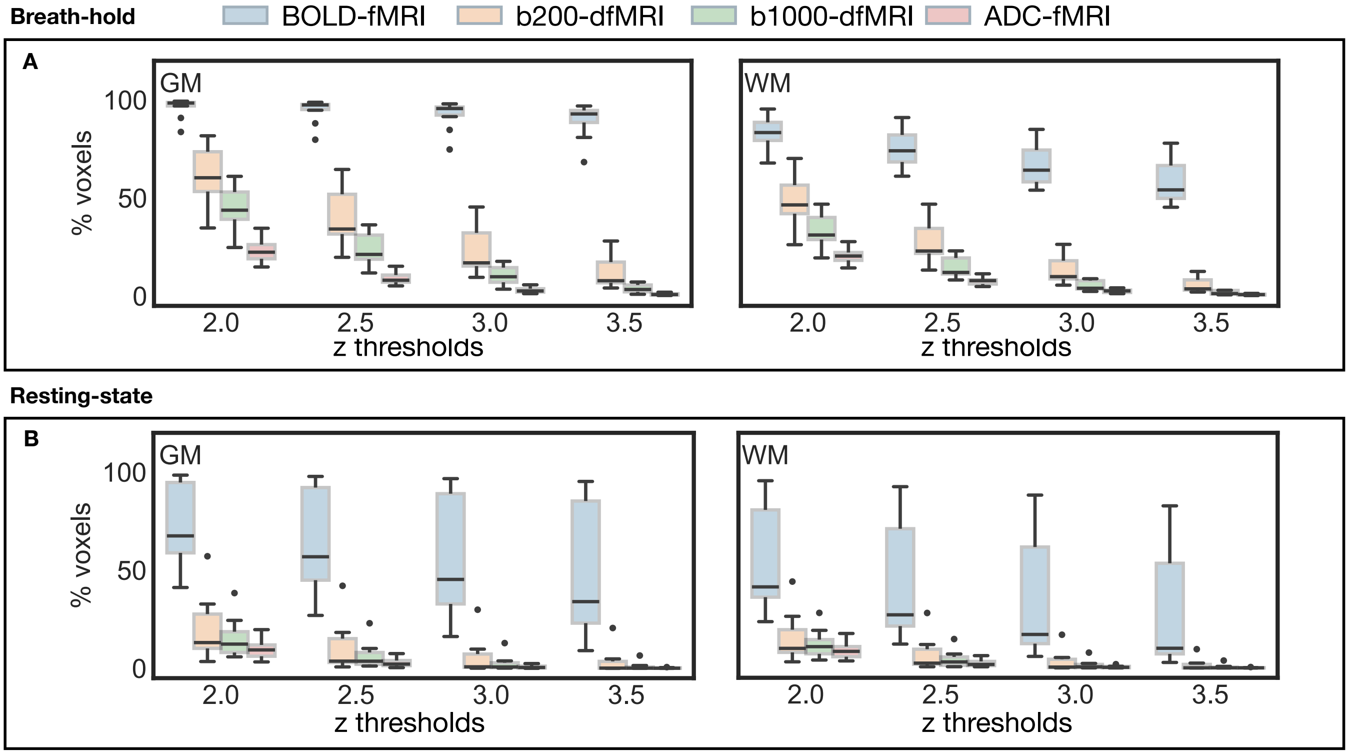


Supplementary Figure 7: **Summary statistics:** Percentage of voxels showing significant association between $p_{ET}CO_2$ and fMRI at different z thresholds. A) Breath-hold: BOLD-fMRI (n = 12, 22 runs), b200-dfMRI, b1000-dfMRI and ADC-fMRI (n = 11, 20 runs). B) Resting-state: BOLD-fMRI (n = 11), b200-dfMRI, b1000-dfMRI and ADC-fMRI (n = 10). The first column shows results in significant grey matter (GM) voxels, while the second column shows results in white matter (WM) voxels.

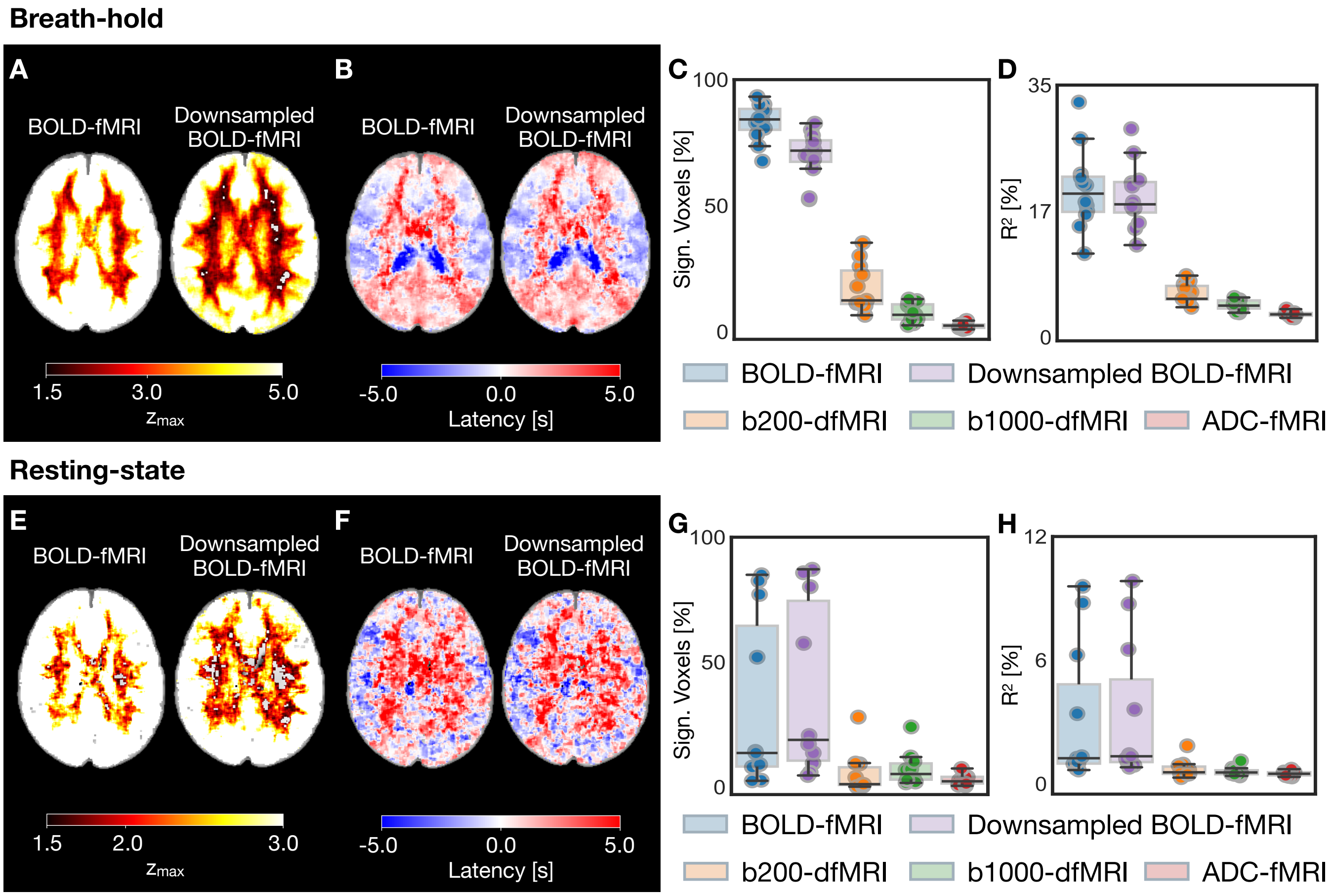


Supplementary Figure 8: **Downsampled BOLD-fMRI: $p_{ET}CO_2$ vs fMRI association** A-D) Breath-hold: BOLD-fMRI (TR = 1 s), downsampled BOLD-fMRI (TR = 2 s) (n = 12, 22 runs), b200-dfMRI, b1000-dfMRI and ADC-fMRI (TR = 2 s)(n = 11, 20 runs). E-H) Resting-state: BOLD-fMRI (TR = 1.1 s), downsampled BOLD-fMRI (TR = 2.2 s) (n = 11), b200-dfMRI, b1000-dfMRI and ADC-fMRI (TR = 2.2 s, n = 10). A) and E) $z_{max}$ maps, B) and F) latency maps for BOLD-fMRI vs downsampled BOLD-fMRI, C) and G) percentage significant voxels, D) and H) coefficient of determination (percentage variance explained) in all voxels. Brain maps show MNI slice z = 95. For breath-hold, the z-threshold used is $z_{max} > 3.09$, for all contrasts. For resting-state, the following contrast-specific z-thresholds, derived from surrogate analysis, were used to determine voxel significance: 4.42 (BOLD-fMRI); 2.95 (Downsampled BOLD-fMRI); 2.84 (b200-dfMRI); 2.36 (b1000- dfMRI); 2.48 (ADC-fMRI).

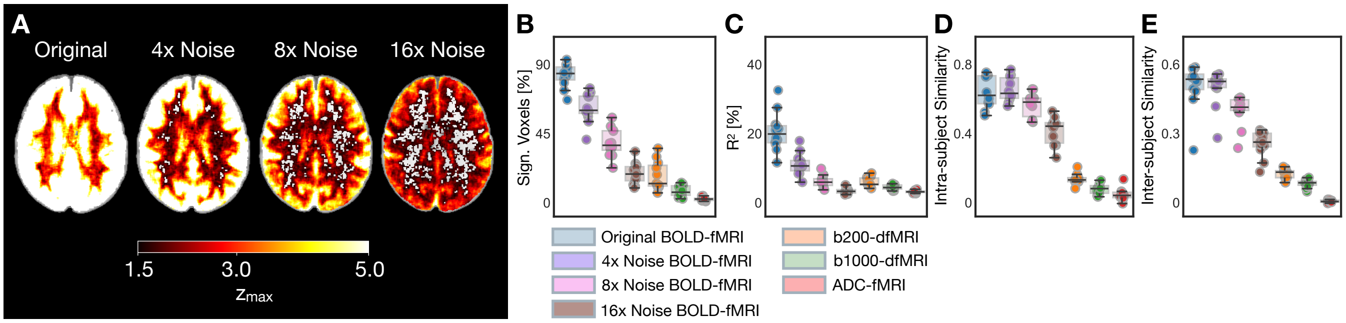


Supplementary Figure 9: **BOLD-fMRI breath-hold with amplified noise levels**. A) Group-average $z_{max}$ maps for the original BOLD-fMRI data, and data with 4x and 8x the noise levels. Slice at $z = 95$ in MNI space is shown. B) Significant voxels ($z_{max} > 3.09$) as a percentage of voxels in the brain. C) Coefficient of determination (percentage variance explained) in all voxels. D) Intra-subject similarity in breath-hold $z_{max}$ maps between task runs. E) Inter-subject similarity in breath-hold $z_{max}$ maps; each point represents the mean similarity from the $z_{max}$ map of one subject (averaged across available runs) to all other subjects.

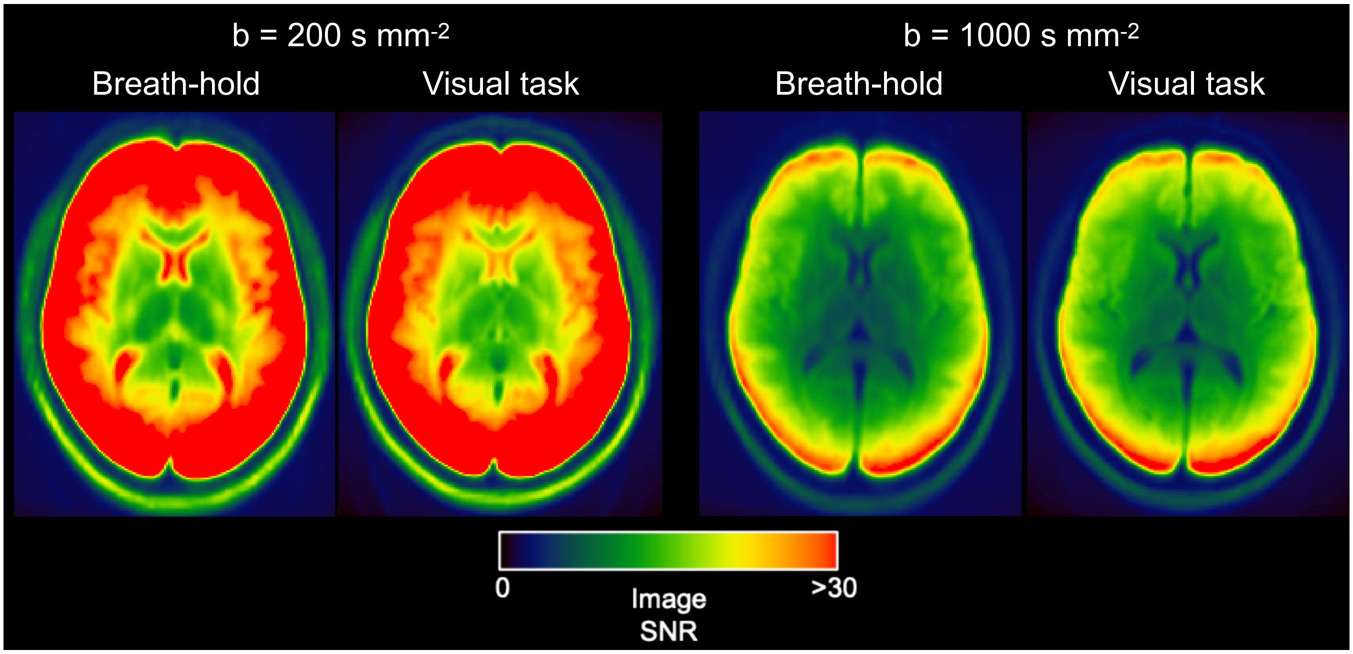


Supplementary Figure 10: **Image SNR comparison** between the breath-hold data used in this study and visual task data from Spencer et al. (2025). The ADC-fMRI acquisition used in this study builds on that of Spencer et al. (2025) with the addition of a cross-term compensated waveform. SNR values are compared in Supplementary Table 3.

### CSF Contribution

We first evaluated the potential CSF contribution to the diffusion-weighted signal using a three-compartment model (Rydhög et al., 2017), which combines the free-water model (Pasternak et al., 2009) with the IVIM model (Le Bihan et al., 1988). Specifically, we employed the following equation:

$$\frac{S}{S_0} = f_{CSF}e^{-b\,ADC_{CSF}} + f_{blood}e^{-b\,ADC_{blood}} + f_{tissue}e^{-b\,ADC_{tissue}}$$

We found that a change in CSF fraction of $\Delta f_{CSF}$ = -0.6% (Donahue et al., 2006; Jin & Kim, 2010) yields an overall ADC change of -0.28%, when calculating initial and final ADC from signals measured at $b_1 = 200$ and $b_2 = 1000$ s mm$^{-2}$ ($f_{csf} = 0.035$; $f_{blood} = 0.05$; $f_{tissue} = 0.915$; $ADC_{CSF}$ = 3 x $10^{-3}$ mm$^2$ s$^{-1}$; $ADC_{blood}$ = 10 x $10^{-3}$ mm$^2$ s$^{-1}$; $ADC_{tissue}$ = 1 x $10^{-3}$ mm$^2$ s$^{-1}$; initial ADC = 1.03614 x $10^{-3}$ mm$^2$ s$^{-1}$; final ADC = 1.03324 x $10^{-3}$ mm$^2$ s$^{-1}$). It also translates into a b200-dfMRI and a b1000-dfMRI signal decrease of -0.32% and -0.09%, respectively. Given that compartment volume fractions differ throughout the brain, we also evaluated a range of initial $f_{csf}$ (1 to 25%, where the variation is compensated by tissue fraction), and a range of $\Delta f_{csf}$ (-0.6 to 0%, where the variation is compensated by blood fraction), which resulted in a ΔADC of at most -0.30%.

Compression of the perivascular space (PVS) may result in a reduction in CSF mobility, and therefore a decrease in $ADC_{CSF}$. While we cannot precisely quantify the $ADC_{CSF}$ decrease associated with PVS compression, a 5% reduction in $ADC_{CSF}$ in the equation above would result in a decrease in measured ADC of less than 0.02%, with $f_{csf}$ up to 0.05 (previously reported PVS volume fractions are in the range 0.01-5%; Pham et al. 2022). Thus the effect of PVS compression on measured ADC is expected to be negligible.

To experimentally quantify possible CSF confounds, we analysed the average functional signal within voxels containing CSF. Specifically, partial volume estimates (pve) of CSF were obtained by applying FSL FAST to the $T_1$-weighted volume. We then averaged the timeseries within three pve ranges (pve = 0, 0 < pve $0.25$, and 0.25 < pve < 0.5). The resulting average timeseries are shown in Supplementary Figure 11. For voxels with pve > 0, BOLD-fMRI, b200-dfMRI, and b1000-dfMRI all exhibited task-related signal increase of 1%, 0.5% and 0.5%, respectively, peaking around 20 s after the breath-hold task onset, whereas the ADC-fMRI signal remained comparatively stable.

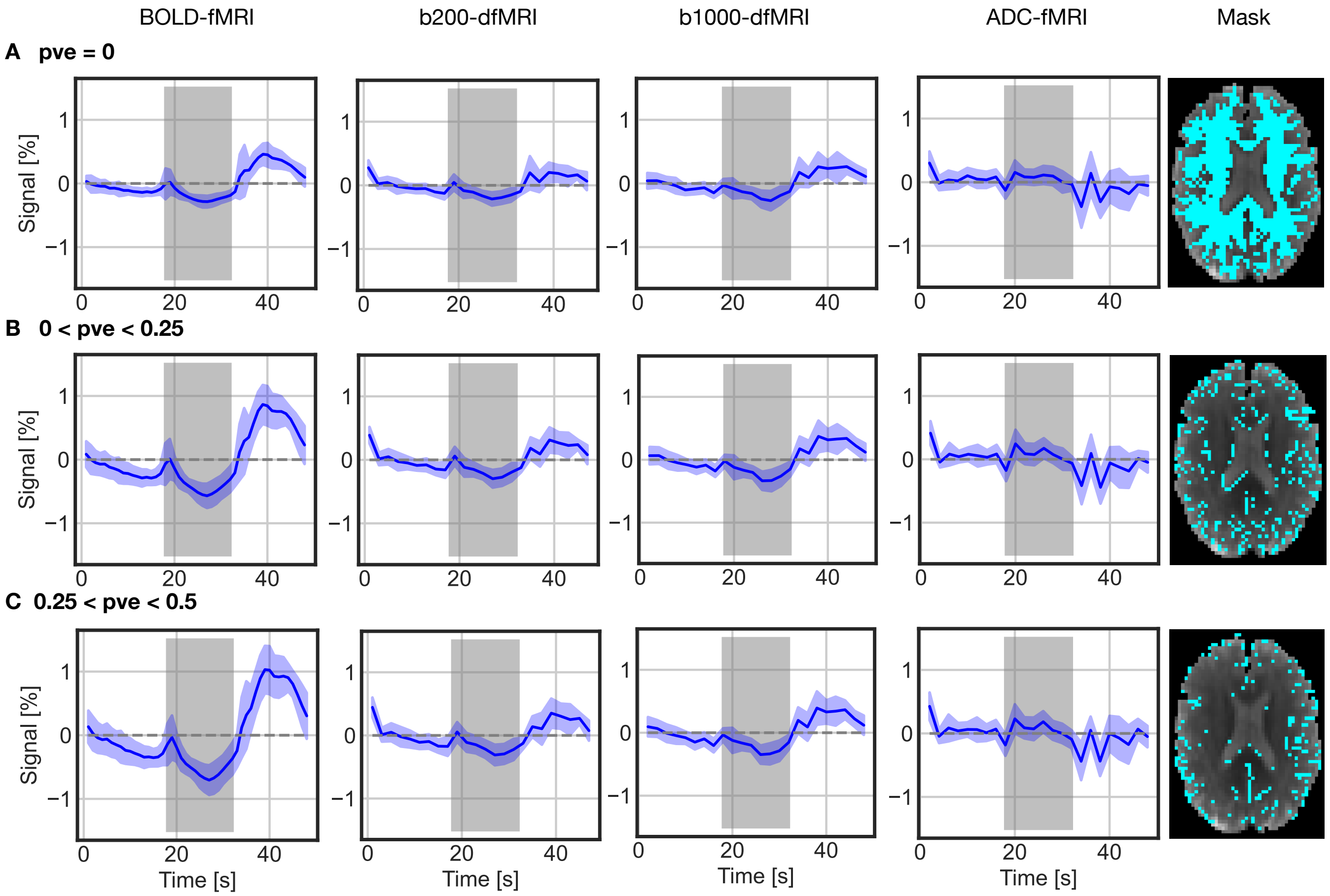


Supplementary Figure 11: **Group-average timeseries in voxels with partial volume with CSF**, in breath-hold task: BOLD-fMRI (n = 12, 22 runs), b200-dfMRI, b1000-dfMRI and ADC-fMRI (n = 11, 20 runs). Partial volume estimate (pve) maps of CSF were generated with FSL fast, and masks corresponding to A) pve = 0, B) 0 < pve < 0.25, and C) 0.25 < pve < 0.5, were applied (displayed in the fifth column, for an example subject). Shaded regions indicate the group standard deviation. It should be noted that the high frequency signal fluctuations aligned with task onset and offset may result from field inhomogeneities caused by pronounced successive chest movements (exhale, inhale) or from head movements occurring in synchrony with the task.

### Resting-State Functional Connectivity

To demonstrate the presence of functional signals in the resting-state ADC-fMRI data, we conducted functional connectivity analysis for each subject. Following previously described preprocessing steps (with the exception of spatial smoothing), further cleaning of physiological noise was applied using independent component analysis (ICA) as follows (Griffanti et al., 2014; McKeown et al., 1998). ICA was performed using FSL's Melodic (Jenkinson et al., 2012) to separate each functional timecourse into 100 components. Noise components were manually classified based on the spatial maps, timecourses and power spectra (Griffanti et al., 2014; Salimi-Khorshidi et al., 2014). Components identified as noise mostly originated from movement, CSF pulsations, or artifacts associated with the multiband acquisition. Noise components were then regressed from the data, along with motion parameters and their derivatives. This was applied to the b200-dfMRI and b1000-dfMRI timecourses separately, before combining these to calculate the cleaned ADC-fMRI timecourse.

The $T_1$-weighted image for each subject was parcellated into regions defined by the Deskian-Killiany atlas (Desikan et al., 2006), using Freesurfer (Fischl, 2012), which were then transformed to functional space. Only regions with at least 10 voxels within the functional imaging volume were retained, leaving 78 regions. The mean ADC timeseries within each region was normalised, and a functional connectivity matrix was calculated for each subject by measuring the Pearson correlation coefficient between region timecourses. The resulting correlation values were Fisher z-transformed to allow averaging across subjects.

The resulting group-average resting-state functional connectivity matrix for ADC-fMRI is shown in Supplementary Figure 12. Multiple connectivity clusters can be observed, with the strongest connectivity seen within and between the frontal and parietal lobes, and within the occipital lobe. Notably, interhemispheric connectivity patterns reflect those within hemispheres.

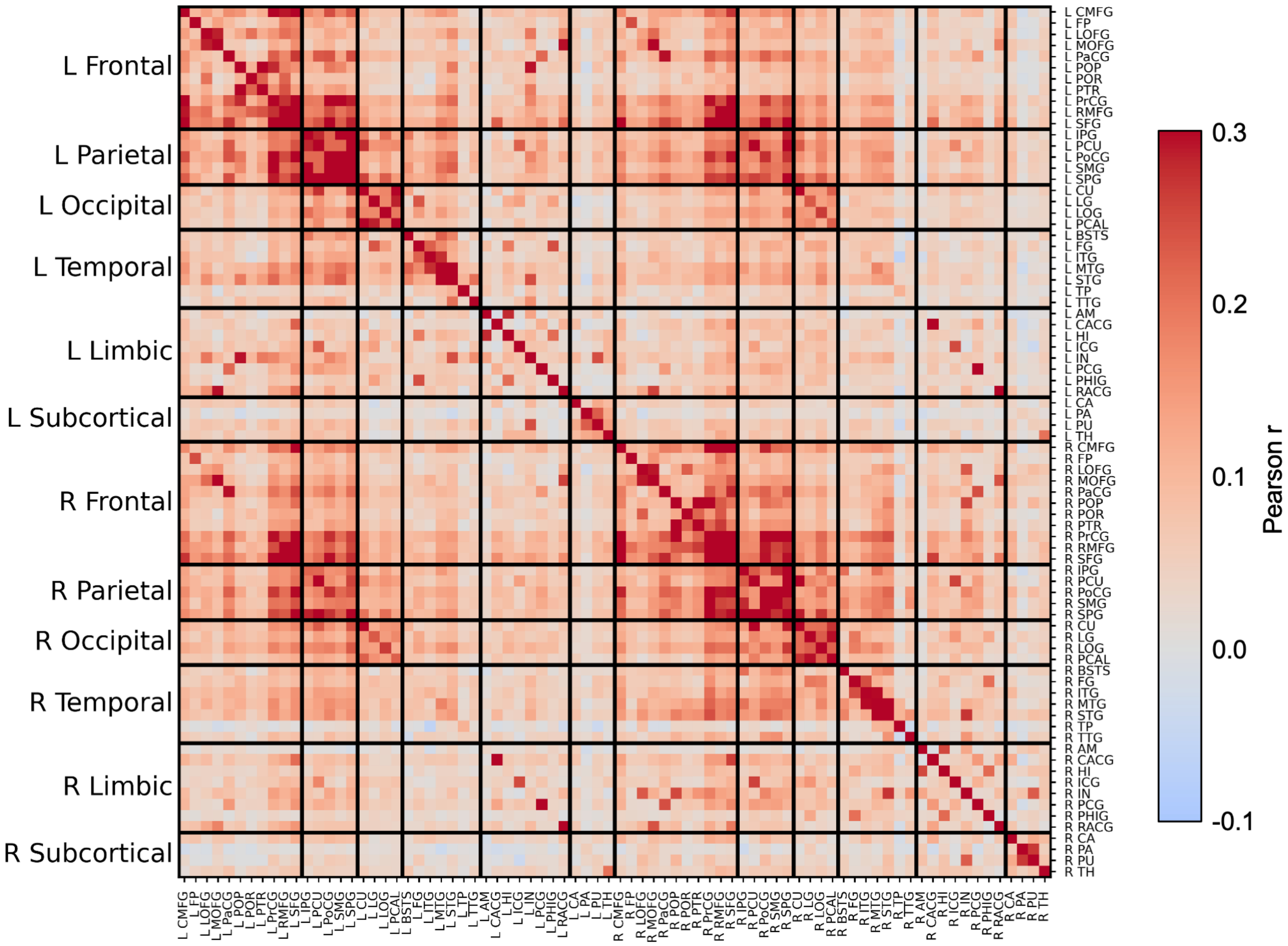


Supplementary Figure 12: **ADC-fMRI resting-state functional connectivity**. Functional connectivity was measured between 78 Desikan-Killiany atlas regions, by measuring the Pearson correlation between region timecourses for each subject. Correlation coefficients were Fisher's z-transformed for averaging between subjects, then group-average values were converted back to Pearson correlation coefficients for plotting.

#### BOLD Denoising

In breath-hold BOLD data, the default TEDICA denoising pipeline was found to remove BOLD components which were associated with the task. These components explained large amounts of variance, with spatial maps covering widespread areas of the brain. Thus, they did not resemble expected BOLD components and were removed by the automatic components classification process. We therefore modified the default decision tree to include the task block design as a regressor, in order to keep any BOLD components which correlated with the breath-hold task.

In resting-state data, the small BOLD signal changes induced by spontaneous fluctuations in resting breathing were not identified as components by TEDICA. As such, the default TEDICA denoising decision tree was used, as this was found to only remove the expected non-BOLD, TE-independent noise components.

In order to further verify that Tedana denoising was not removing respiration signals from the resting-state data, we repeated the cross-correlation analysis between $p_{ET}CO_2$ and resting-state BOLD-fMRI data without denoising. As shown in Supplementary Figure 13, without ICA denoising the variance explained by $p_{ET}CO_2$ was similar to (or slightly lower than) the original results presented in the main text. Thus, Tedana denoising was not removing signals associated with pETCO2.

Note that due to this modification to the breath-hold denoising pipelines, the preprocessing pipelines for resting-state and breath-hold are not exactly the same (see Supplementary Figure 1). As such, we have not directly compared the absolute z-scores or latencies between breath-hold and resting-state in any of our analysis or interpretation. Notably, for the analysis of similarity between breath-hold and resting-state z-score maps, we used a spatial correlation in order to assess whether the spatial patterns of the maps are similar, not the absolute values. For all other results, we have focused on interpretation of the differences between contrasts and patterns across the brain.

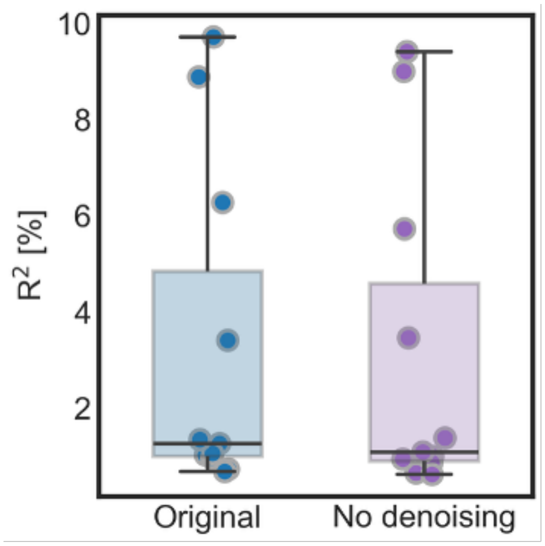


Supplementary Figure 13: **Association between resting-state BOLD data and respiration with and without Tedana denoising**. Coefficient of determination (percentage variance explained) in all voxels for the cross-correlation between contrast and $p_{ET}CO_2$, with and without default Tedana ICA denoising.
